# Molecular pathology of a cystic fibrosis mutation is explained by loss of a hydrogen bond

**DOI:** 10.1101/2021.10.19.464938

**Authors:** Márton A. Simon, László Csanády

**Affiliations:** HCEMM-SE Molecular Channelopathies Research Group, Semmelweis University, Tuzolto u. 37-47, Budapest, H-1094, Hungary; MTA-SE Ion Channel Research Group, Semmelweis University, Tuzolto u. 37-47, Budapest, H-1094, Hungary; Department of Biochemistry, Semmelweis University, Tuzolto u. 37-47, Budapest, H-1094, Hungary

**Author notes:** Corresponding author: László Csanády, M.D., Ph.D., Semmelweis University, Department of Biochemistry, Tuzolto u. 37-47, Budapest, H-1094, Hungary, /60048.

**Keywords:** CFTR, ABC protein, Class III mutant, R117H, gating defect

## Abstract

The phosphorylation-activated anion channel CFTR is gated by an ATP hydrolysis cycle at its two cytosolic nucleotide binding domains, and is essential for epithelial salt-water transport. A large number of CFTR mutations cause cystic fibrosis. Since recent breakthrough in targeted pharmacotherapy, CFTR mutants with impaired gating are candidates for stimulation by potentiator drugs. Thus, understanding the molecular pathology of individual mutations has become important. The relatively common R117H mutation affects an extracellular loop, but nevertheless causes a strong gating defect. Here we identify a hydrogen bond between the side chain of arginine 117 and the backbone carbonyl group of glutamate 1124 in the cryo-electronmicroscopic structure of phosphorylated, ATP-bound CFTR. We address the functional relevance of that interaction for CFTR gating using macroscopic and microscopic inside-out patch-clamp recordings. Employing thermodynamic double-mutant cycles, we systematically track gating-state dependent changes in the strength of the R117-E1124 interaction. We find that the H-bond is formed only in the open state, but neither in the short-lived “flickery” nor in the long-lived “interburst” closed state. Loss of this H-bond explains the entire gating phenotype of the R117H mutant, including robustly shortened burst durations and strongly reduced intraburst open probability. The findings may help targeted potentiator design.

## Introduction

Loss-of-function mutations of the Cystic Fibrosis Transmembrane Conductance Regulator (CFTR) anion channel disrupt transepithelial salt-water transport in the lung, gut, intestine, pancreatic duct, and sweat duct, and cause cystic fibrosis (CF), the most common inherited lethal disease among caucasians (O’Sullivan and Freedman, 2009). Based on their molecular consequences the ∼1700 identified CF mutations have been categorized into several classes, such as those that impair synthesis of the full-length CFTR polypeptide (Class I), processing and trafficking of the CFTR potein (Class II), channel gating (Class III), or anion permeation through the open pore (Class IV) (De Boeck and Amaral, 2016). Until recently symptomatic therapy remained the only available treatment option for CF patients, but this has been profoundly changed by a major breakthrough in the development of small-molecule drugs that target the CFTR protein itself. Treatment by several “corrector” drugs that improve processing and maturation of Class II mutant CFTR protein, and by the “potentiator” compound Vx-770 (ivacaftor) which increases open probability (P_o_) of Class III mutant channels, have lead to significant symptomatic improvement for patients with Class II/III mutations (Ramsey et al., 2011; Davies et al., 2018), and has been approved by the FDA. Because the responsiveness to potentiators of different Class III mutants is greatly variable (Van Goor et al., 2014), understanding the molecular pathologies of such mutations bears strong clinical relevance.

CFTR is an ATP Binding Cassette (ABC) protein which contains two transmembrane domains (TMD1, 2; Fig. 1A, *gray*) and two cytosolic nucleotide binding domains (NBD1, 2; Fig. 1A, *blue* and *green*). These two ABC-typical NBD-TMD halves are linked by a cytosolic R domain (Fig. 1A, *magenta*) which is unique to CFTR and contains multiple serines that must be phosphorylated by cAMP-dependent protein kinase (PKA) to allow channel activity (Riordan et al., 1989). CFTR pore opening/closure (gating) is linked to an ATP hydrolysis cycle at the NBDs and resembles the active transport cycle of ABC exporters (reviewed, e.g., in (Csanády et al., 2019)). In the presence of ATP single phosphorylated CFTR channels open into “bursts”: groups of openings separated by brief (∼10 ms) “flickery” closures and flanked by long (∼1 s) “interburst” closures. Upon ATP binding CFTR’s NBDs associate (Fig. 1B-C) into a tight head-to-tail dimer which is accompanied by a large rearrangement of its TMDs from an inward-facing (Fig. 1B) to an outward-facing (Fig. 1C) orientation. The inward-facing TMD conformation with separated NBDs corresponds to the stable interburst (IB) closed state in which the channel gate, near the extracellular membrane surface, is closed (Liu et al., 2017). The outward-facing TMD conformation, with a tight NBD dimer that occludes two ATP molecules in interfacial binding sites (sites 1 and 2), corresponds to the bursting (B) state in which a large lateral gap between TM helices 4 and 6 continues to connect the pore to the cytosol, bypassing the constricted ABC-transporter inner gate (Zhang et al., 2018b). The B state is a compound state that includes the fully open (O) and the flickery closed (C_f_) states. Although in the outward-facing CFTR structure (pdbid: 6msm) the external end of the pore is not quite as wide as would be required for passage of chloride ions, at present that structure is the best available model for the O state (Zhang et al., 2018b). In contrast, the structure of the C_f_ state is unknown but, based on its fast kinetics, the O↔C_f_ transition is likely a localized movement. For wild-type (WT) CFTR the majority of sojourns in the highly stable B state are terminated by hydrolysis of the ATP at site 2, which prompts NBD dimer dissociation and resets the channel into the inward-facing IB state (Csanády et al., 2010).

**Figure 1.**
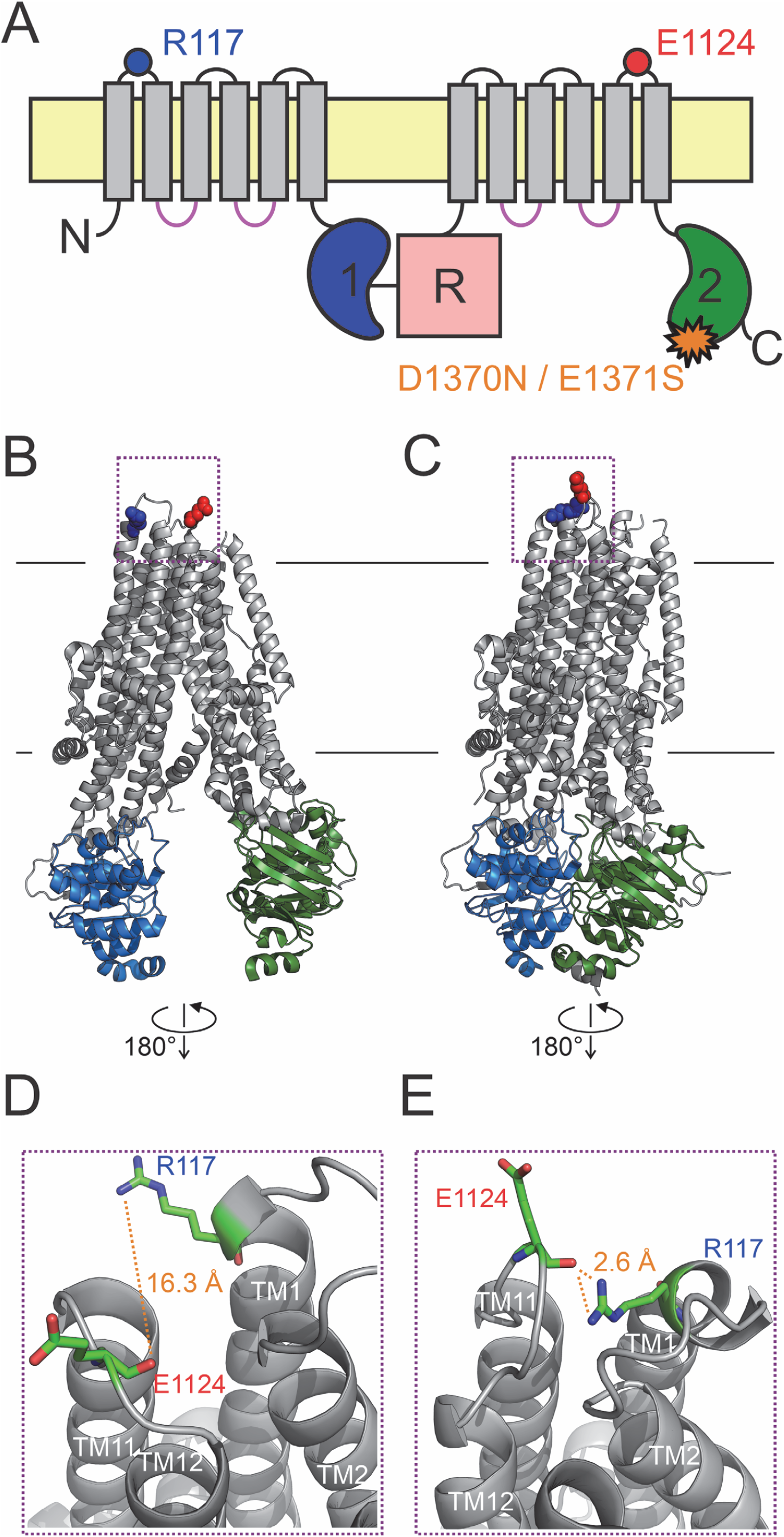
A H-bond between the R117 side chain and the E1124 backbone carbonyl group is apparent in the quasi-open CFTR structure. *A*, Cartoon topology of CFTR and location of target positions (*blue* and *red dot*). TMDs, *gray*; NBD1, *blue*; NBD2, *green*; R domain, *magenta*; membrane, *yellow. Orange star* denotes catalytic site mutation in the NBD2 Walker B motif. *B*-*C*, Target residues R117 (*blue*) and E1124 (*red*) shown in space fill on the cryo-EM structures of (*B*) inward-facing dephospho-CFTR (pdbid: 5uak) and (*C*) outward-facing phosphorylated ATP-bound CFTR (pdbid: 6msm). Domain color coding as in *A*, the R domain is not depicted. *D*-*E*, Close-up views of the extracellular region highlighted in *B*-*C* by *dotted purple boxes*. Target residues are shown as *sticks*, target distances are illustrated by *orange dotted lines*.

Substitution of the conserved arginine at position 117, and in particular mutation R117H, is among the most common CF mutations ((Dean et al., 1990; Wilschanski et al., 1995); https://cftr2.org/). In heterologous expression systems the R117H mutation does not impair surface expression (Sheppard et al., 1993; Hammerle et al., 2001), but severely reduces whole-cell CFTR currents (Sheppard et al., 1993). Consistent with its location in the first extracellular loop (ECL1), replacement of the R117 side chain (Fig. 1A, *blue dot*) causes a mild conductance defect suggesting that its positive charge is normally involved in recruiting extracellular chloride ions to the outer mouth of the pore (Sheppard et al., 1993; Zhou et al., 2008). However, the robust reduction of whole-cell currents is not explained by that small conductance defect, but rather by a strong gating defect (Sheppard et al., 1993) which places the R117H mutation into Class III. The reduced P_o_ of the mutant is primarily caused by a large reduction in the mean burst duration (τ_burst_) (Sheppard et al., 1993; Yu et al., 2016). In addition, the kinetics of intraburst gating is also altered in the mutant: shortened open times are paired with prolonged flickery closed times (Yu et al., 2016). Because R117H-CFTR currents are only modestly stimulated by Vx-770 (Van Goor et al., 2014; Yu et al., 2016), understanding the molecular pathology of this mutation will be important for developing better drugs optimized for personalized treatment.

How a mutation at the outer mouth of the pore may affect channel gating, thought to be controlled by NBD dimer formation/disruption, seemed puzzling at first. CFTR’s cyclic gating mechanism precludes equilibrium thermodynamic analyses (Csanády, 2009), and makes it difficult to discern whether a mutational effect on burst durations reflects an alteration in the rate of ATP hydrolysis or in the stability of the prehydrolytic open state. Using a non-hydrolytic site 2 mutation which reduces CFTR gating in saturating ATP to reversible IB↔B transitions, Φ-value analysis (Auerbach, 2007) revealed that the pore opening (IB→B) step is a spreading conformational wave initiated at site 2 of the NBD dimer interface and propagated along the longitudinal protein axis towards the extracellular surface. In the high-energy transition state (T^‡^) site 2 has already adopted its B-state conformation (i.e., it is already tightly dimerized) whereas the extracellular portion of the TMDs, which includes the gate, is still in its IB-state (closed) conformation (Sorum et al., 2015). Thus, mutations at extracellular positions, like 117, are expected to similarly perturb the stabilities of the T^‡^ state and the IB closed state, and thus to selectively impact closing rate but not opening rate. That expectation was later confirmed for a series of perturbations at position 117: because Q, A, H, and C substitutions here all selectively accelerated closure (shortened τ_burst_), it was concluded that in wild-type CFTR the R117 side chain is involved in a stabilizing interaction that is formed only in the B state but neither in the IB state nor in the T^‡^ state for opening (Sorum et al., 2017). However, in the absence of high-resolution structures the interaction partner of the R117 side chain could not be identified.

Here we exploit the recent cryo-electronmicroscopic (cryo-EM) structures of human CFTR obtained in inward- and outward-facing conformations (Liu et al., 2017; Zhang et al., 2018b), to identify the functionally important molecular interaction partner of the R117 side chain. We then test the functional role of this putative interaction using detailed single-channel kinetic analysis, and employ thermodynamic double mutant cycles to rigorously quantitate how its strength changes between specific gating states. Our analysis provides a mechanistic explanation for the strong energetic role that the conserved arginine at position 117 plays in CFTR channel gating, as well as for the molecular pathology caused by its mutations. It also provides new structural inference for the organization of the C_f_ state.

## Results

### Cryo-EM structures suggest a strong H-bond between the R117 side chain and the E1124 peptide carbonyl group in the open-channel state

In cryo-EM structures of both inward-facing (Fig. 1B) ((Liu et al., 2017); pdbid: 5uak) and outward-facing (Fig. 1C) ((Zhang et al., 2018b); pdbid: 6msm) CFTR the R117 side chain is clearly resolved (Fig. 1B-C, *blue space fill*). In the quasi-open structure it approaches, and forms a strong H-bond with, the backbone carbonyl oxygen of residue E1124 (Fig. 1E), located in ECL6 (Fig. 1A, *red dot*). In the closed-pore structure it is pointing out towards the solvent, and is not in contact with any other residue of the protein (Fig. 1D). The latter structure does not represent the closed state of an active channel gating in the presence of ATP, as it was obtained from unphosphorylated CFTR in the absence of ATP, and correspondingly contains density for the unphosphorylated R domain wedged in between the two unliganded NBDs and the cytosolic ends of the TMD helices. Nevertheless, functional studies suggest that the overall organization of the TMDs in this structure resembles that of the “active” closed state (Bai et al., 2011; Cui et al., 2014; Wang et al., 2014). If so, then the identified 117-1124 H-bond might represent an interaction that changes dynamically during the gating cycle. To study the functional relevance of this putative interaction for CFTR gating energetics, we employed two different background mutations which disrupt ATP hydrolysis at site 2 (Fig. 1A, *orange star*), and reduce CFTR gating in saturating ATP to a reversible IB↔B mechanism.

### Deletion of residue 1124 reproduces the R117 phenotype

The NBD2 Walker B glutamate (E1371) side chain forms the catalytic base in site 2, and its mutations disrupt ATP hydrolysis (Zhang et al., 2018b). Correspondingly, the E1371S mutation prolongs τ_burst_ by >100-fold (Vergani et al., 2003; Csanády et al., 2013), revealing slow reversibility of the IB→B step. That slow rate of the (non-hydrolytic) B→IB transition is conveniently measured in macroscopic inside-out patch recordings by activating pre-phosphorylated E1371S CFTR channels using a brief exposure to 5 mM ATP, and then observing the slow rate of macroscopic current decay upon sudden ATP removal (Fig. 2A, *black trace*); the time constant of a fitted exponential reports τ_burst_ (Fig. 2B, *black bar*), i.e., the average life time of the B state. Introduction of the R117H mutation into this background robustly accelerated the current decay (Fig. 2A, *blue trace*), yielding ∼6-fold shortening of τ_burst_ (Fig. 2B, *blue bar*), consistent with an earlier report using the E1371Q background (Yu et al., 2016).

**Figure 2.**
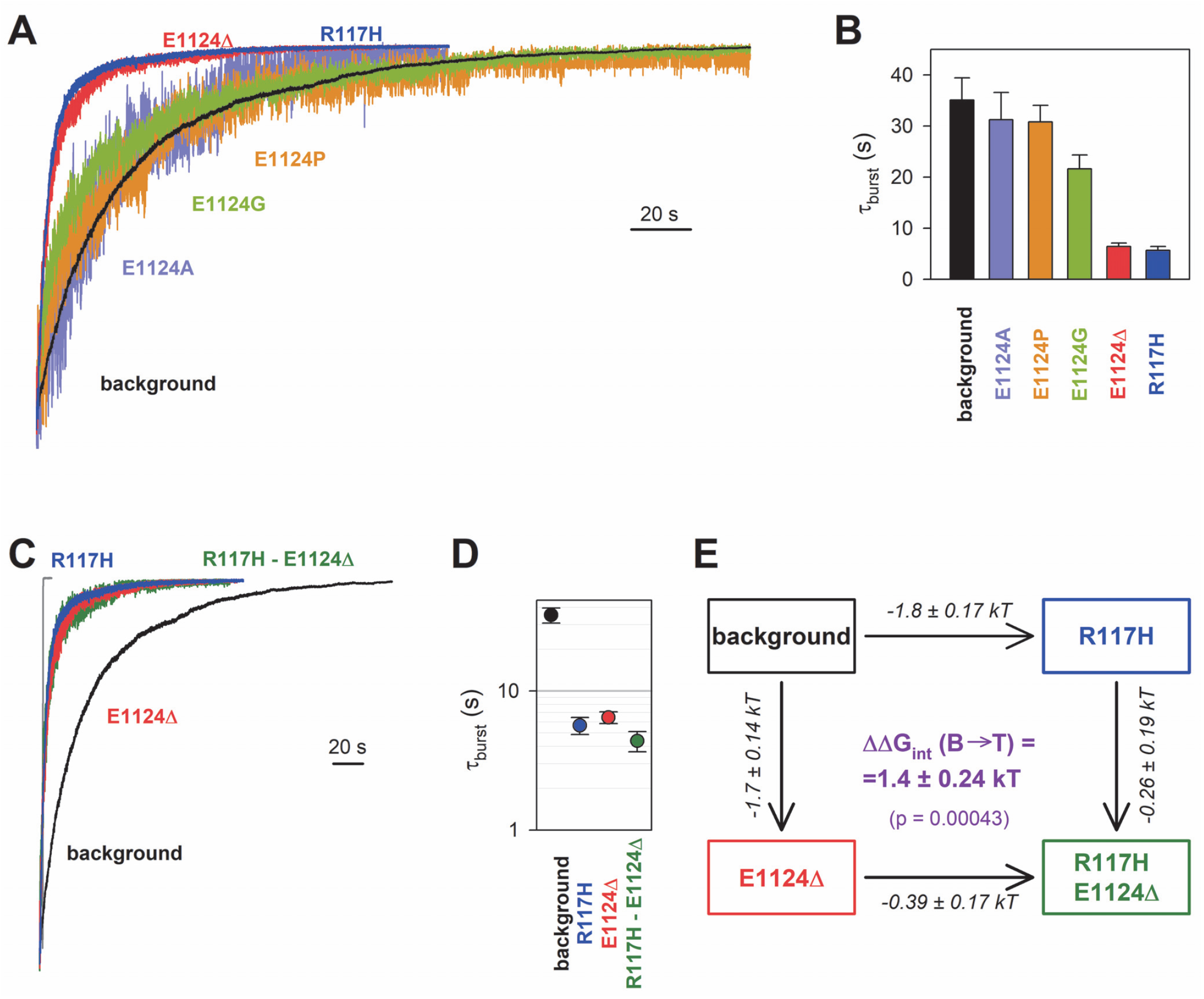
Energetic coupling between positions 117 and 1124 changes during non-hydrolytic closure. *A*-*C*, Macroscopic current relaxations following ATP removal for indicated CFTR mutants (*color coded*) harboring the E1371S background mutation. Inside-out patch currents were activated by exposure of pre-phosphorylated channels to 5 mM ATP. Current amplitudes are shown rescaled by their steady-state values. The *gray trace* in *C* illustrates the speed of solution exchange, estimated as the time constant of deactivation (∼100 ms) upon Ca^2+^ removal of the endogenous Ca^2+^-activated choride current, evoked by brief exposure to Ca^2+^. Membrane potential (V_m_) was -20 mV. *B, D*, Relaxation time constants of the currents in *A* and *C*, respectively, obtained by fits to single exponentials. Data are shown as mean±S.E.M. from 6-10 experiments. Note logarithmic ordinate in *D. E*, Thermodynamic mutant cycle showing mutation-induced changes in the height of the free enthalpy barrier for the B→IB transition (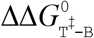, *numbers on arrows*; *k*, Boltzmann’s constant, *T*, absolute temperature). Each corner is represented by the mutations introduced into positions 117 and 1124 of E1371S-CFTR. 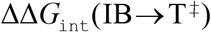 (*purple number*) is obtained as difference between 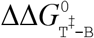 values along two parallel sides of the cycle.

If the phenotype caused by the R117H mutation is indeed due to loss of the R117-E1124 interaction, then a similar phenotype should be brought about by perturbations of position 1124. However, designing a suitable perturbation here is complicated by the fact that E1124 interacts through its peptide carbonyl oxygen. Notably, positioning of a backbone carbonyl group is unaffected by side chain substitutions, and its targeted elimination using nonsense suppression-based unnatural amino acid incorporation (Pless and Ahern, 2013) has not yet been developed. As expected, truncation of the E1124 side chain using the E1124A mutation did not affect closing rate (Fig. 2A, *violet trace*; Fig. 2B, *violet bar*), confirming that the E1124 side chain is not required for normal gating. A slight shortening of τ_burst_ by the E1124G mutation suggested that increasing flexibility of ECL6 only marginally accelerates closing rate (Fig. 2A, *green trace*; Fig. 2B, *green bar*). We next examined whether insertion of a proline might distort ECL6 sufficiently to move the E1124 backbone carbonyl group out of reach of the R117 side chain. However, neither the E1124P (Fig. 2A, *orange trace*; Fig. 2B, *orange bar*) nor the E1126P (Fig. 2 – Fig. Suppl. 1) mutation caused a shortening of τ_burst_. Thus, the only viable strategy to increase the R117-E1124 distance appeared to be shortening of the ECL6 loop using a single-residue deletion. To obtain an estimate of the expectable distance increase we resorted to *in silico* modelling. Using the quasi-open CFTR structure as a starting point, molecular dynamics simulations (see Material and Methods) were used to estimate the mean 117-1124 bond distance in WT CFTR, as well as following *in silico* single-residue substitutions or deletions around position 1124 (Fig. 2 – Fig. Suppl. 2). These experiments confirmed unchanged bond distances for the proline mutants, and suggested deletion of residue 1124 itself as the only simple way to increase the distance between the R117 side-chain guanidino group and the nearest ECL6 backbone carbonyl (in this case that of G1123). Indeed, in our E1371S background construct introduction of the E1124Δ mutation produced a phenotype very similar to that of R117H, accelerating the macroscopic current decay (Fig. 2A, *red trace*), and shortening τ_burst_ by ∼6-fold (Fig. 2B, *red bar*).

### Energetic coupling between positions 117 and 1124 changes during non-hydrolytic closure

Functional effects of mutations of any individual amino acid residue in a protein provide little information regarding the role of a specific hypothesized residue-residue interaction, because in the WT protein the targeted residue might be involved in multiple interactions, all of which are altered by its mutations. To dissect and quantify the energetic effect of perturbing a specific residue-residue interaction, thermodynamic mutant cycle analysis must be employed (Vergani et al., 2005). The concept behind that approach is that the background construct, two single mutants that each perturb one or the other target residue, and the corresponding double mutant form a thermodynamic cycle. If the two target residues do not interact, or if their interaction remains static (does not change between various gating states), then the energetic effects of mutating either target residue will be additive in the double mutant. Conversely, if the strength of the interaction between the target residues changes between gating states S_1_ and S_2_, then the effect on the relative stability of state S_2_ caused by mutating one target residue 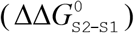 will depend on the nature of the residue present at the other target position – thus, the energetic effects of the two target-site mutations will not be additive in the double mutant. The change in strength of the target interaction during step S_1_→S_2_ (ΔΔ*G*_int_ (S_1_→ S_2_)) is obtained as the difference between 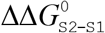 values along two parallel sides of the mutant cycle (Fig.2E).

Introducing the R117H mutation into an E1124Δ background caused only marginal further acceleration of E1371S non-hydrolytic closing rate (Fig. 2C, *green trace*), shortening τ_burst_ ∼1.47-fold (Fig. 2D, *green* vs. *red symbol*). In energetic terms, the ∼6-fold shortening of τ_burst_ brought about by the R117H mutation alone (Fig. 2D, *blue* vs. *black symbol*) signals an ∼1.8 *kT* decrease in the height of the free enthalpy barrier for the B→IB transition (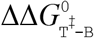 Fig. 2E, number on *top horizontal arrow*). In contrast, the ∼1.47-fold acceleration caused by the same mutation in the E1124Δ background reports a decrease in the same barrier height by only ∼0.39 *kT* (Fig. 2E, number on *bottom horizontal arrow*). The difference between those two numbers quantifies the change in the strength of the R117-E1124 interaction in an E1371S channel while it progresses from the B state to the T^‡^ state, on its way towards the IB closed state (Fig. 2E, *purple number*). That value of 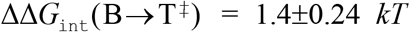 is significantly different from zero (p=0.00043), and its positive signature reports that a stabilizing interaction present in the B state is lost in the T^‡^ state. (The theoretical alternative of a lack of interaction in the B state but a destabilizing interaction in the T^‡^ state is inconsistent with structural evidence.)

### The stabilizing interaction between the target positions is present only in the bursting state

To address how the 117-1124 interaction changes between the IB state and the T^‡^ state, mutational effects on opening rate need to be quantified in steady-state single-channel recordings. Because gating of the E1371S background construct is prohibitively slow for steady-state single-channel analysis, we employed a different non-hydrolytic background mutation, D1370N, which removes the Walker B glutamate side chain that coordinates catalytic Mg^2+^ in site 2 (Hung et al., 1998; Rai et al., 2006; Gunderson and Kopito, 1995). This mutation disrupts ATP hydrolysis but results in shorter burst durations, allowing steady-state kinetic analysis of the IB↔B gating process (Csanády et al., 2010; Sorum et al., 2015). The R117H, E1124Δ, and R117H/E11214Δ target site mutations were introduced into this background, and gating of single prephosphorylated channels was studied in the presence of 5 mM ATP (Fig. 3A), a saturating concentration for all four constructs (Fig. 3 – Fig. Suppl. 1). In such steady-state recordings flickery and interburst closures can be readily discriminated by dwell-time analysis (Materials and Methods; Fig. 3 – Fig. Suppl. 2), allowing calculation of not only τ_burst_, but also of mean IB durations (τ_interburst_) (Table 1).

**Table 1.** Model-independent descriptive gating parameters for the indicated mutant CFTR constructs in the D1370N background. Gating parameters were obtained from steady-state recordings as described in Materials and Methods. Data are displayed as mean±S.E.M. from 5-8 patches.

| | $\tau_{\text{burst}}$ (ms) | $\tau_{\text{interburst}}$ (ms) | $\tau_{\text{flicker}}$ (ms) | $n_{\text{flicker}}$ | $P_o$ |
| --- | --- | --- | --- | --- | --- |
| **background (D1370N)** | $2000 \pm 250$ | $5100 \pm 1100$ | $10 \pm 1.6$ | $4.8 \pm 0.82$ | $0.35 \pm 0.056$ |
| **R117H** | $67 \pm 8.8$ | $2800 \pm 800$ | $33 \pm 5.0$ | $1.2 \pm 0.12$ | $0.017 \pm 0.0055$ |
| **E1124Δ** | $100 \pm 16$ | $4800 \pm 810$ | $23 \pm 4.0$ | $2.1 \pm 0.41$ | $0.013 \pm 0.0033$ |
| **R117H**<br>**E1124Δ** | $58 \pm 4.4$ | $3200 \pm 880$ | $50 \pm 7.9$ | $1.1 \pm 0.099$ | $0.0054 \pm 0.0011$ |

**Figure 3.**
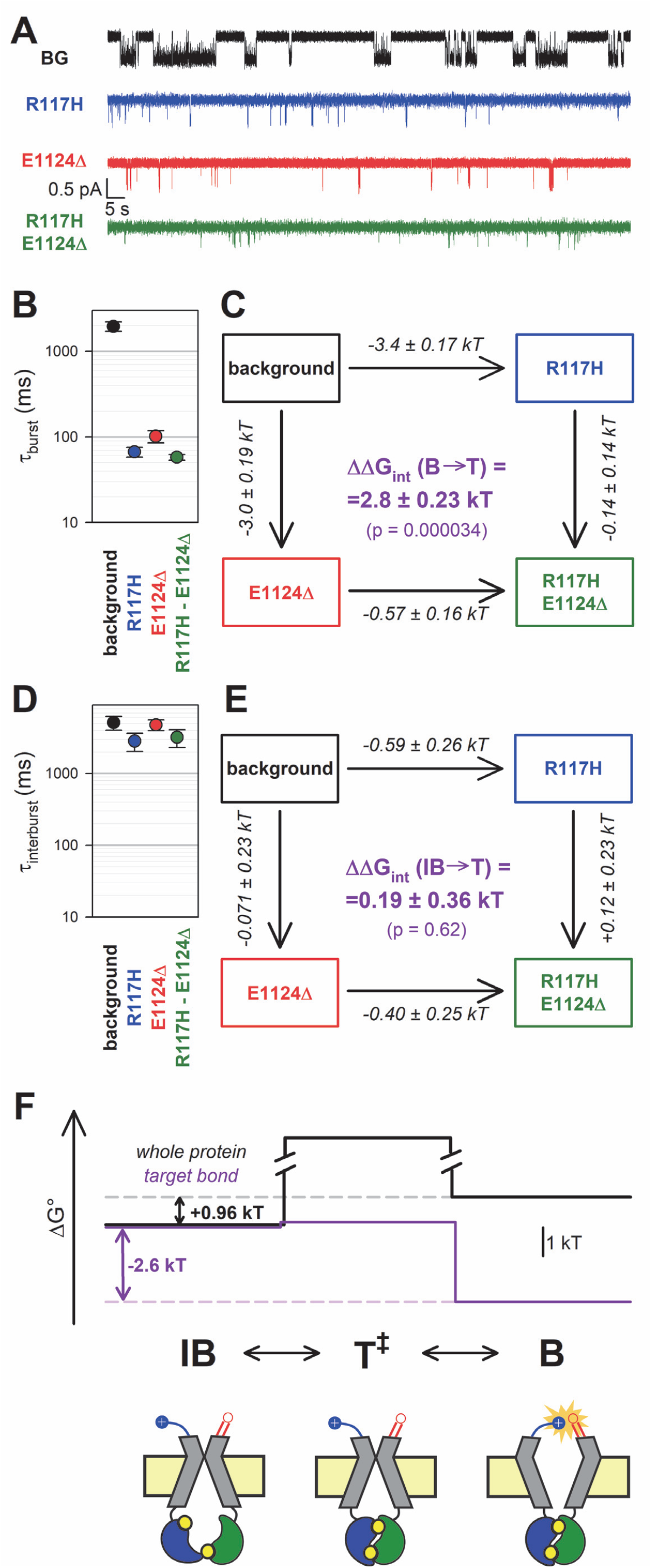
The H-bond between the target positions is formed only in the bursting state. *A*, Single-channel currents in 5 mM ATP of indicated CFTR mutants (*color coded*) harboring the D1370N background (BG) mutation. Channels were pre-phosphorylated by ∼1-minute exposure to 300 nM PKA + 5 mM ATP. V_m_ was -80 mV. *B, D*, Mean burst (*B*) and interburst (*D*) durations obtained by steady-state dwell-time analysis (see Materials and Methods). Data are shown as mean±S.E.M. from 5-8 patches; note logarithmic ordinates. *C, E*, Thermodynamic mutant cycles showing mutation-induced changes in the height of the free enthalpy barrier for the (*C*) B→IB and (*E*) IB→B transition (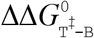 and 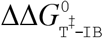, respectively, *numbers on arrows*; *k*, Boltzmann’s constant, *T*, absolute temperature) Each corner is represented by the mutations introduced into positions 117 and 1124 of D1370N-CFTR. ΔΔ*G*_int_ (*purple number*) is obtained as difference between ΔΔ*G*^0^ values along two parallel sides of the cycle. *F*, (*Top*) Free enthalpy profile of the entire channel protein (*black curve*) and of the 117-1124 interaction (*purple curve*) along the states of the IB↔B gating process (see Materials and Methods). (*Bottom*) Cartoon representation of channel conformations in the IB, T^‡^, and B states. *Color coding* as in Fig. 1A; ATP, *yellow circles. Blue* and *red* moieties represent the R117 side chain and the E1124 backbone carbonyl group, respectively. *Yellow star* in state B represents formation of the H-bond.

Mutational effects on closing rate resembled those observed in the E1371S background: both the R117H and the E1124Δ mutation robustly shortened τ_burst_ (by ∼30-fold and ∼20-fold, respectively, Fig. 3B, *blue* and *red* symbols, respectively, vs. *black symbol*), whereas little further shortening was observed in the double mutant (Fig. 3B, *green symbol*). Thus, whereas the ∼30-fold shortening of τ_burst_ by the R117H mutation alone (Fig. 3B, *blue* vs. *black symbol*) reports a decrease in 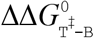 by ∼3.4 *kT* (Fig. 3C, *top horizontal arrow*), the ∼1.76-fold shortening of τ_burst_ by the same mutation in the E1124Δ background (Fig. 3B, *green* vs. *red symbol*) reports a decrease in the height of the same barrier by only ∼0.57 *kT* (Fig. 3C, *bottom horizontal arrow*). The disparity between those two values, 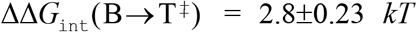 (Fig. 3C, *purple number*), is significantly different from zero (p=0.000034) confirming disruption of a strong stabilizing interaction between the target positions during the B→T^‡^ transition.

In contrast, channel opening rates were little affected by the target site mutations, with τ_interburst_ values remaining within twofold for all four constructs (Fig. 3D, Table 1). Correspondingly, the calculated change in the strength of the 117-1124 interaction during step IB→T^‡^, 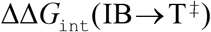 (Fig. 3E, *purple number*), was not significantly different from zero. Of note, correct estimation of τ_interburst_ strongly depends on correct estimation of the number of channels in the patch, which becomes increasingly difficult when the P_o_ is very low. Thus, for our target-site mutants and the double mutant the presence of more than one channel in the patch was excluded by strongly boosting channel P_o_ at the end of each recording using N^6^-(2-phenylethyl)-dATP (P-dATP) and/or the potentiator drug Vx-770 (Fig. 3 – Fig. Suppl. 3, also see Materials and Methods).

Taken together, these steady-state data reveal that in D1370N channels a strong stabilizing interaction between positions 117 and 1124 is formed in state B, but is absent in states T^‡^ and IB (Fig. 3F, *purple* bond energy profile; Fig. 3F *bottom*, cartoon).

### The H-bond between the target positions is broken in the flickery closed state

The bursting state comprises the open (O) and flickery closed (C_f_) states. If the strength of an interaction changes between states O and C_f_ then perturbing that interaction should affect intraburst P_o_ (P_O|B_), i.e., the fraction of time the pore is open within a burst. To address whether the R117-E1124 H-bond is disrupted in the flickery closed state, we studied additivity of effects of the R117H and E1124Δ mutations on the intraburst closed-open equilibrium constant K_eq|B_, obtained as K_eq|B_ = P_O|B_/(1-P_O|B_). Effects on intraburst gating kinetics are most conveniently studied in the E1371S background, by briefly applying ATP to activate the channels and then focusing on the last bursting channel following ATP removal (Fig. 4A, (Csanády et al., 2010; Yu et al., 2016)). Because in the absence of bath ATP reopening from the IB state is no longer possible, in such “last-channel” segments of record all closures (except for the final) are necessarily flickery closures, and the long life time of the B state in E1371S allows sampling of a large number of O↔C_f_ transitions.

**Figure 4.**
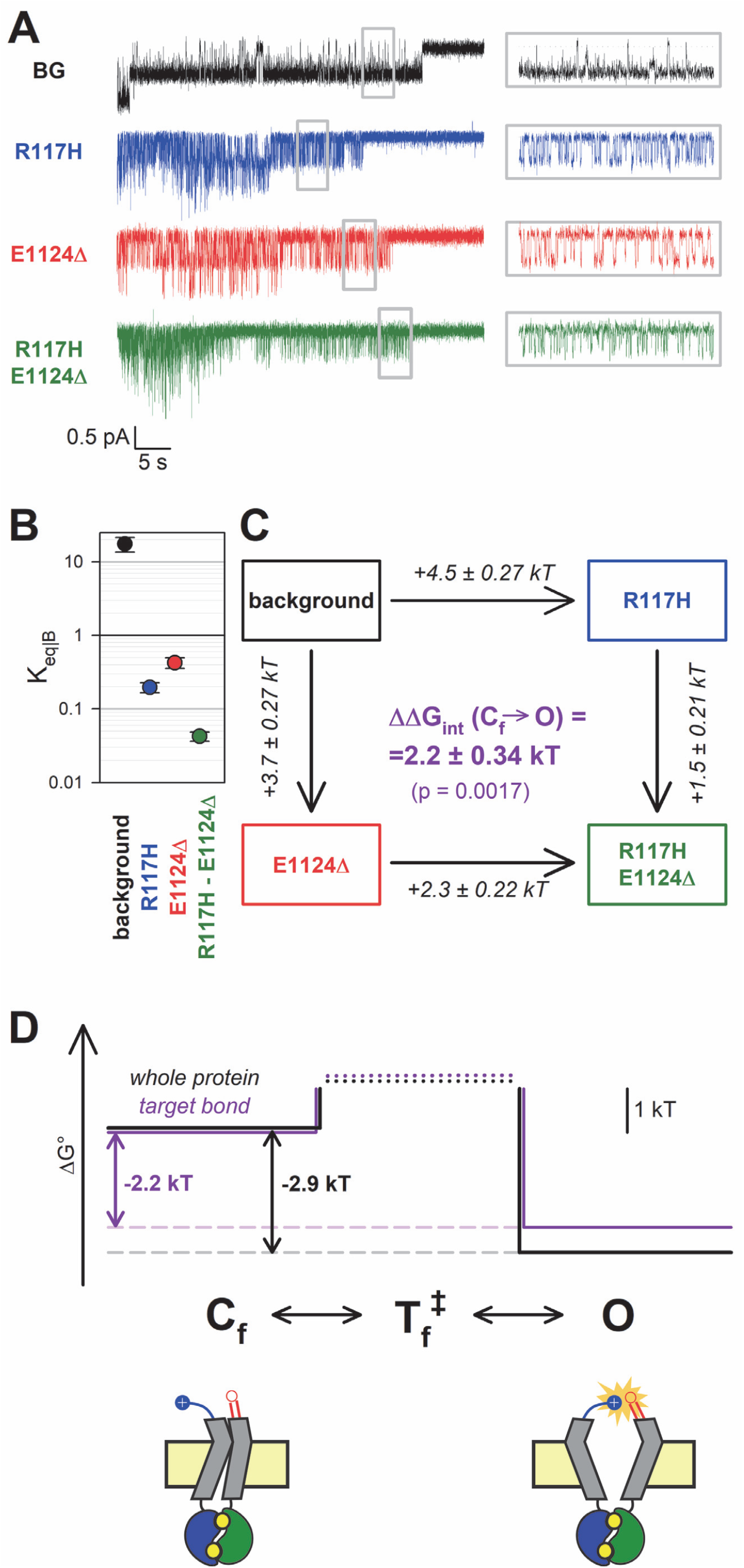
The H-bond between the target positions is broken in the flickery closed state. *A*, Currents of last open channels surviving ATP removal, for indicated CFTR mutants (*color coded*) harboring the E1371S background mutation. Inside-out patch currents were activated by exposure of pre-phosphorylated channels to 5 mM ATP. Insets (*right*) show 5-second segments corresponding to *gray boxes* at an expanded time scale. V_m_ was -80 mV. *B*, Closed-open equilibrium constants within a burst (see Materials and Methods). Data are shown as mean±S.E.M. from 5-6 patches; note logarithmic ordinate. *C* Thermodynamic mutant cycle showing mutation-induced changes in the stability of the O state relative to the C_f_ state (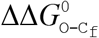, *numbers on arrows*; *k*, Boltzmann’s constant, *T*, absolute temperature). Each corner is represented by the mutations introduced into positions 117 and 1124 of E1371S-CFTR. ΔΔ*G*_int_ (*purple number*) is obtained as the difference between ΔΔ*G*^0^ values along two parallel sides of the cycle. *D*, (*Top*) Free enthalpy profile of the entire channel protein (*black curve*) and of the 117-1124 interaction (*purple curve*) along the states of the intraburst (O↔C_f_) gating process (see Materials and Methods). (*Bottom*) Cartoon representation of channel conformations in the C_f_ and O states. *Color coding* as in Fig. 3F.

As reported previously (Yu et al., 2016), the R117H mutation caused a strong reduction in P_O|B_, due to an ∼20-fold shortening of mean open times (τ_open_) and a 5-fold prolongation of mean flickery closed times (τ_flicker_) (Fig. 4A, *insets, blue* vs. *black trace*; Table 2). Interestingly, this aspect of the R117H phenotype was also reproduced by the E1124Δ mutant, which demonstrated a similarly lowered P_O|B_ (Fig. 4A, *red trace*, Table 2). Although P_O|B_ was even further reduced for R117H-E1124Δ (Fig. 4A, *green trace*, Table 2), the effects of the two single mutations on K_eq|B_ were not additive in the double mutant. Whereas the ∼90-fold reduction in K_eq|B_ caused by the R117H mutation alone (Fig. 4B, *blue* vs. *black symbol*) reported an increase in the C_f_-O free enthalpy difference 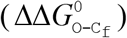 by ∼4.5 *kT* (Fig. 4C, *top vertical arrow*), the same mutation reduced K_eq|B_ only ∼10-fold (Fig. 4B, *green* vs. *red symbol*), amounting to 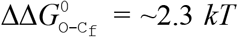 (Fig. 4C, *bottom vertical arrow*), when introduced into an E1124Δ background. The obtained interaction energy, ΔΔ*G*_int_ (C_f_ →O) = 2.2±0.34 *kT* (Fig. 4C, *purple number*), is significantly different from zero (p=0.0017) indicating that, even within the bursting state, the R117-E1124 H-bond is formed only in the open state, not in the flickery closed state (Fig. 4D, *purple* bond energy profile; Fig. 4D *bottom*, cartoon).

**Table 2.** Model-independent descriptive parameters of intraburst gating for the indicated mutant CFTR constructs in the E1371S background. Gating parameters were obtained from last-open channel recordings as described in Materials and Methods. Data are displayed as mean±S.E.M. from 5-6 patches.

| | $\tau_{\text{flicker}}$ (ms) | $\tau_{\text{open}}$ (ms) | $K_{\text{eq\vert B}}$ |
| --- | --- | --- | --- |
| **background (E1371S)** | $11 \pm 2.3$ | $200 \pm 27$ | $17 \pm 3.9$ |
| **R117H** | $58 \pm 7.3$ | $11 \pm 1.1$ | $0.20 \pm 0.031$ |
| **E1124Δ** | $42 \pm 6.0$ | $18 \pm 1.7$ | $0.42 \pm 0.070$ |
| **R117H**<br>**E1124Δ** | $150 \pm 18$ | $6.2 \pm 0.50$ | $0.042 \pm 0.0060$ |

### Both kinetic schemes used to describe CFTR bursting behavior adequately explain observed effects

CFTR’s bursting behaviour may be explained by two alternative linear three-state models, C_s(low)_↔C_f(ast)_↔O or C_s_↔O↔C_f_. Because those two models cannot be differentiated in any individual steady-state recording, the true model is still unknown. All the analysis presented so far was therefore performed in a model-independent manner. Nevertheless, we also sought to address whether the mutational effects observed here might provide any support for one or the other kinetic mechanism.

Maximum likelihood fits (Csanády, 2000) by the C_s_↔C_f_↔O and C_s_↔O↔C_f_ model of steady-state single-channel recordings, obtained for the four constructs in the D1370N background (Fig. 3A), yielded rate constants for all four microscopic transition rates of the respective scheme (Figs. 5C and 5D, *center*, average rates (in s^-1^) are *color coded* by mutation). Thus, for both models, a mutant cycle could be built on each of the four transition rates, to quantitate ΔΔ*G*_int_ of the target H-bond for the respective gating step (Fig. 5 – Fig. Suppl. 1). Finally, those four ΔΔ*G*_int_ values allowed reconstruction of the entire 117-1124 interaction free enthalpy profile assuming one or the other model (Fig. 5A-B, *purple plots*). Assuming either model, these profiles predict formation of the target H-bond selectively in state O. Correspondingly, a comparison of the standard free enthalpy profiles of gating for each of the four constructs (Fig. 5C-D, *top*), calculated for both models from the fitted microscopic transition rate constants (see Materials and Methods), unanimously predict selective destabilization of the O state by the target site mutations (Fig. 5C-D, *top*, colored profiles vs. *black profile*). Moreover, using the C_s_↔O↔C_f_ model the change in target bond strength between states O and C_s_ (∼2.6 *kT*) and between states O and C_f_ (∼2.5 *kT*) turned out to be nearly identical (Fig. 5A). All in all, the data are equally consistent with either model.

**Figure 5.**
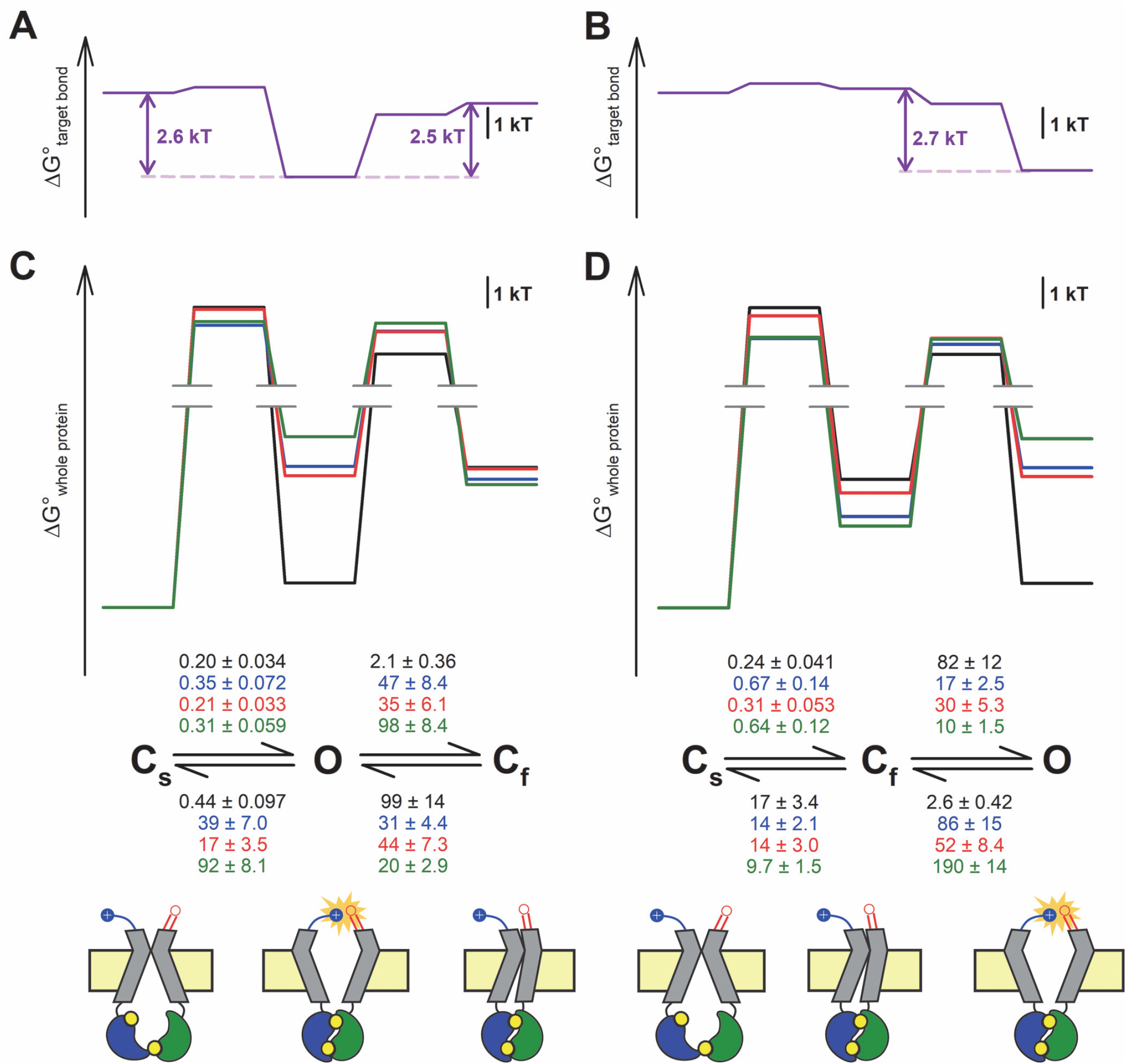
Both alternative linear three-state models adequately explain the effects of the mutations. *A-D*, Free enthalpy profiles (*A*-*B*) of the 117-1124 interaction in D1370N-CFTR, and (*C*-*D*) of the entire channel protein for various mutants (*color coded*) in the D1370N background, along all states of the gating process assuming either the C_s_↔O↔C_f_ (*A, C*) or the C_s_↔C_f_↔O (*B, D*) model. The profiles in (*A*-*B*) were obtained from the ΔΔ*G*_int_ values for the four gating steps, those in (*C*-*D*) from the microscopic transition rate constants (numbers in s^-1^, *color coded* by mutants) obtained by fitting either model to the steady-state recordings in Fig. 3A (see Materials and Methods). Cartoons in *C*-*D* (bottom) represent the channel conformations in the C_s_, O, and C_f_ states. *Color coding* as in Fig. 3F.

## Discussion

Here we have shown that a strong H-bond between the side chain of R117 and the backbone carbonyl group of E1124 is formed in the open state but neither in the interburst nor in the flickery closed state of the CFTR channel. We have further shown that loss of that hydrogen bond is responsible for the entire gating phenotype caused by CF mutation R117H, including a dramatic shortening of burst durations and a strong reduction of intraburst P_o_. These findings have clarified the molecular pathology of a relatively common CF mutation (and of the less frequent variants R117C/G/L/P), and might prove useful for targeted design of better potentiator drugs for this mutant.

The fact that it is the backbone carbonyl group, not the side chain, of E1124 that participates in this H-bond is consistent with several observations. First, mutations of position 1124 have not been identified in CF patients, consistent with our conclusion that the E1124 side chain does not play an important role. Second, whereas the arginine at the position corresponding to 117 in hsCFTR is absolutely conserved across species, the residue at the position equivalent to 1124 is not (Fig. 2, Fig. Suppl. 3). On the other hand, the *lengths* of both ECL6 and ECL1 are absolutely conserved (Fig. 2, Fig. Suppl. 3), consistent with 3D spacing between those two loops playing an important role.

Based on homology modeling the R117 side chain was earlier proposed to interact with the E1126 side chain (Cui et al., 2014). Our data suggest that is not the case, consistent with cryo-EM structures of both quasi-open and closed CFTR in which the E1126 side chain points away from R117. Indeed, mutation E1126P which truncates the E1124 side chain did not result in an R117H-like phenotype, but rather prolonged τ_burst_ (Fig. 2 – Fig. Suppl. 1). The reason for the latter effect is unclear, but maybe due to changes in flexibility of the ECL6 loop which natively contains three glycines (sequence: GEGEG; Fig. 2 – Fig. Suppl. 3). Mutation E1126P (GEG<u>E</u>G → GEG<u>P</u>G) is expected to reduce, whereas mutation E1124G (G<u>E</u>GEG → G<u>G</u>GEG) is expected to increase ECL6 flexibility: correspondigly, τ_burst_ was prolonged by the former, but slightly shortened by the latter (Fig. 2A-B, *green*) mutation.

Based on the structures the IB→B transition involves a large conformational rearrangement during which dozens of intramolecular interactions are broken and dozens of others are formed (Liu et al., 2017; Zhang et al., 2018b). How can disruption of a single H-bond result in such a pronounced gating phenotype as that caused by the R117H mutationã Although both conformations of the protein are stabilized by large numbers of local interactions, evolution has carefully balanced the energetic contributions of these, such that net Δ*G*^0^ for the overall IB→B protein conformational change is not far from zero. Indeed, for the D1370N background construct that net Δ*G*^0^ for the entire protein (∼1 *kT*; Fig. 3F, *black free enthalpy profile*) is smaller than the bond energy of our single target H-bond (∼2.6 *kT*; Fig. 3F, *purple free enthalpy profile*). Thus, just as the balance of an equilibrated two-armed scale is thrown off by removal of a single small weight from any of its two trays, no matter how heavily both trays had been loaded, elimination of a single H-bond selectively present in the B state can profoundly shift the gating equilibrium towards the IB conformation.

In contrast, based on its fast kinetics, the O↔C_f_ transition may involve a smaller conformational change with only a few local interactions broken/formed. Because the overall Δ*G*^0^ for that transition is again comparable to that of formation/disruption of the 117-1124 H-bond (∼2.9 *kT* vs. ∼2.2 *kT* in the E1371S background construct; Fig. 4D, *black* vs. *purple free enthalpy profile*), that bond might represent a key interaction that governs intraburst gating in CFTR. In the cryo-EM structure of phosphorylated ATP-bound zebrafish CFTR ((Zhang et al., 2017); pdbid: 5w81), although the NBDs are dimerized and the TMDs outward-facing, the external extremity of the pore is tightly sealed. The latter feature is due to a localized displacement of the external portions of TM helices 1 and 12, as compared to the quasi-open structure of human CFTR. Interestingly, that displacement increases the ECL1-ECL6 distance in the zebrafish structure, moving the residues equivalent to our target positions (R118 and D1132) too far away to form an H-bond. Thus, out of two suggested possible alternative explanations (Zhang et al., 2017) our findings support the interpretation (Zhang et al., 2018a) that the zebrafish outward-facing structure might represent the flickery closed state (cf., cartooned C_f_ state in Figs. 4-5).

The magnitude of the effect on B-state stability of disrupting the 117-1124 H-bond depended on the choice of the non-hydrolytic model: τ_burst_ was shortened by ∼6-fold in the E1371S (Fig. 2C-D), but by ∼30-fold in the D1370N (Fig. 3A-B) background. That discrepancy suggests that the precise arrangement of the ECLs in the B state is not exactly identical in those two constructs, so that the distance between the two target positions, and thus the strength of the H-bond between them, is slightly different (1.4 *kT* (Fig. 2E) vs. 2.8 *kT* (Fig. 3C)). Which of those two values is more representative of a WT channelã Considering the effect of the R117H mutation on τ_burst_ of WT CFTR a semi-quantitative lower estimate can be made. In an earlier study (Yu et al., 2016) the rate of the B→IB transition was increased from ∼3.5 s^-1^ in WT CFTR to ∼11 s^-1^ in R117H CFTR. Since the life time of the posthydrolytic B state is very short (Csanády et al., 2010), that rate essentially reflects the life time of the prehydrolytic B state, and is approximated by the sum of the rates for ATP hydrolysis (*k*_h_) and for non-hydrolytic closure (*k*_nh_). In WT channels hydrolytic closure dominates (*k*_nh_<<*k*_h_; (Csanády et al., 2010)), i.e., *k*_h_>3 s^-1^, *k*_nh_<0.5 s^-1^ (cf., *k*_nh_∼0.5 s^-1^ for the fastest-closing among several known non-hydrolytic models, D1370N (Fig. 3B, *black symbol*)). Based on the plausible assumption that the ECL1 mutation R117H does not alter the catalytic rate in site 2 of the NBD dimer, in R117H CFTR *k*_h_ remains ∼3 s^-1^, and thus *k*_nh_ is ∼8 s^-1^. These arguments provide a lowest possible estimate of ∼16-fold for the increase in non-hydrolytic closing rate caused by the R117H mutation in WT CFTR, more in line with our findings in the D1370N background.

CFTR bursting may be accounted for by two possible linear three state models, C_s_↔O↔C_f_ and C_s_↔C_f_↔O. In the past, multiple studies have addressed whether ATP-dependence (Winter et al., 1994), voltage dependence (Cai et al., 2003), or pH-dependence (Chen et al., 2017) of CFTR gating might allow differentiation between those two mechanisms, but in each case both models proved equally adequate to explain the data. Because the 117-1124 H-bond is formed only in state O but not in states C_s_ or C_f_ (Figs. 3-4), we reasoned that mutant cycles built on microscopic transition rates obtained by fits to one or the other model might provide some support for a choice. In particular, if analysis based on the C_s_↔O↔C_f_ model had returned significantly different values of ΔΔ*G*_int_ for the C_s_↔O and C_f_↔O steps, that would have rendered this model highly unlikely, given that in both of those steps ΔΔ*G*_int_ should reflect the strength of the same H-bond. However, no such discrepancy emerged from our analysis (Fig. 5A), thus, our data are adequately explained by either model. Moreover, there is currently no structural evidence to support the existence of a fixed order of transitions among the three channel conformations seen in the cryo-EM structures of dephosphorylated human, phosphorylated ATP-bound zebrafish, and phosphorylated ATP-bound human CFTR, that are thought to resemble states C_s_, C_f_, and O, respectively (cartooned in Fig. 5C, bottom). Thus, more structural information will be required to finally settle this question.

## Materials and Methods

### Molecular biology

Mutations were introduced into the human CFTR(E1371S)/pGEMHE and CFTR(D1370N)/pGEMHE sequences using the QuikChange II XL Kit (Agilent Technologies) and confirmed by automated sequencing (LGC Genomics GmbH). To generate cRNA, the cDNA was linearized (Nhe I HF, New England Biolabs) and transcribed *in vitro* using T7 polymerase (Thermo Fisher mMessage mMachine T7 Kit), cRNA was stored at -80 °C.

### Functional expression of human CFTR constructs in Xenopus laevis oocytes

Oocytes were removed from anaesthetized *Xenopus laevis* following Institutional Animal Care Committee guidelines, and separated by collagenase treatment (Gibco, Collagenase type II). Isolated oocytes were kept at 18°C, in a modified frog Ringer’s solution (in mM: 82 NaCl, 2 KCl, 1 MgCl_2_, and 5 HEPES, pH 7.5 with NaOH) supplemented with 1.8 mM CaCl_2_ and 50 μg/ml gentamycin. Injections by 0.1-10 ng of cRNA, to obtain microscopic or macroscopic currents, was done in a fixed 50 nl volume (Drummond Nanoject II). Recordings were performed 1-3 days after injection.

### Excised inside-out patch recording

The patch pipette solution contained (in mM): 138 NMDG, 2 MgCl_2_, 5 HEPES, pH=7.4 with HCl. The bath solution contained (in mM): 138 NMDG, 2 MgCl_2_, 5 HEPES, 0.5 EGTA, pH=7.1 with HCl. Following excision into the inside-out configuration patches were moved into a flow chamber in which the composition of the continuously flowing bath solution could be exchanged with a time constant of <100 ms using electronic valves (ALA-VM8, Ala Scientific Instruments). CFTR channel gating was studied at 25°C, in the presence of 5 mM MgATP (Sigma), following activation by ∼1 minute exposure to 300 nM bovine PKA catalytic subunit (Sigma P2645). Macroscopic currents were recorded at -20 mV, microscopic currents at -80 mV membrane potential. For the low-P_o_ mutants R117H, E1124Δ, and R117H-E1124Δ evaluation of the number of active channels in the patch was facilitated by stimulating the channels with 50 μM N^6^-(2-phenylethyl)-dATP (P-dATP; Biolog LSI), with or without 10 nM Vx-770 (Selleck Chemicals), at the end of each experiment. Currents were amplified and low-pass filtered at 1 kHz (Axopatch 200B, Molecular Devices), digitized at a sampling rate of 10 kHz (Digidata 1550B, Molecular Devices) and recorded to disk (Pclamp 11, Molecular Devices).

### Kinetic analysis of electrophysiological data

To obtain relaxation time constants (Fig. 2B, D), macroscopic current relaxations were fitted to single exponentials by non-linear least squares (Clampfit 11).

For kinetic analysis of steady-state single-channel recordings in the D1370N background, baseline-subtracted unitary currents, from recordings with no superimposed channel openings, were Gaussian-filtered at 100 Hz and idealized by half-amplitude threshold crossing. Recordings were further analyzed only if the presence of a second active channel could be excluded with >90% confidence using statistical tests described earlier (Csanády et al., 2000). All events lists were fitted with both the C_1(s)_↔O_3_↔C_2(f)_ and the C_1(s)_↔C_2(f)_↔O_3_ model using a maximum likelihood approach which accounted for the presence of an imposed fixed dead time of 4 ms, and returned a set of microscopic transition rate constants *k*_12_, *k*_21_, *k*_23_, and *k*_32_ for both models (Csanády, 2000). The descriptive kinetic parameters τ_burst_, τ_interburst_, τ_flicker_, and *n*_flicker_ (average number of flickery closures per burst) were calculated from the rate constants as follows (Colquhoun and Sigworth, 1995). For the C_1_↔O_3_↔C_2_ model τ_burst_=(1/*k*_31_)(1+*k*_32_/*k*_23_), τ_interburst_=1/*k*_13_, τ_flicker_=1/*k*_23_, *n*_flicker_=*k*_32_/*k*_31_. For the C_1_↔C_2_↔O_3_ model τ_burst_=(1/*k*_21_)((*k*_21_+*k*_23_)/*k*_32_+*k*_23_/(*k*_21_+*k*_23_)), τ_interburst_=((*k*_12_+*k*_21_+*k*_23_)/(*k*_12_*k*_23_))+1/(*k*_21_+*k*_23_), τ_flicker_=1/(*k*_21_+*k*_23_), *n*_flicker_=*k*_23_/*k*_21_. Importantly, fits using either model necessarily yield identical sets of calculated parameters τ_burst_, τ_interburst_, τ_flicker_, and *n*_flicker_, thus, the latter descriptive parameters (Fig. 3B-D, Table 1) are model-independent. The rate of the IB→B step was taken as 1/τ_interburst_, and that of the B→IB step as 1/τ_burst_,

For intraburst kinetic analysis of the last open channel in the E1371S background, mean open times (τ_open_) and mean flickery closed times (τ_flicker_) were obtained as the simple arithmetic averages of the mean open and closed dwell-time durations, respectively, and *K*_eq|B_ was calculated as *K*_eq|B_=τ_open_/τ_flicker_ (Fig. 4B, Table 2).

### Mutant cycle analysis

Changes in the strength of the R117-E1124 interaction between various gating states were evaluated using thermodynamic mutant cycle analysis as described (Mihályi et al., 2016). For an S_1_↔T^‡^↔S_2_ gating step mutation-induced changes in the relative stabilities of those three states were calculated as follows. The change in the height of a transition-state barrier, e.g., for step S_1_→T^‡^, was calculated as 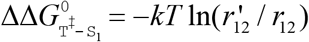, where *k* is Boltzmann’s constant, *T* is absolute temperature, and *r*_12_ and 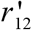 are the rates for the S_1_→S_2_ transition in the background construct and in the mutant, respectively. The change in the stability of the two ground states relative to each other was calculated as 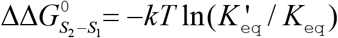, where *K*_eq_ and 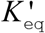 are the equilibrium constants for the S_1_↔S_2_ transition in the background construct and in the mutant, respectively. Interaction free energy (ΔΔ*G*_int_) was defined as the difference between ΔΔ*G*^0^ values along two parallel sides of a mutant cycle. All ΔΔ*G* values are given as mean±S.E.M; S.E.M. values were estimated assuming that *r*_ij_ and *K*_eq_ are normally distributed random variables, using second-order approximations of the exact integrals (Mihályi et al., 2016).

### Calculation of free enthalpy profiles

To construct standard free enthalpy profiles for channel gating (Figs. 3F, 4D, *black profiles*; Fig. 5C-D) ΔΔ*G*^0^ between two ground states was calculated as ΔΔ*G*^0^ = −*kT* ln*K*_eq_. The absolute values of the barrier heights were not calculated (see breaks), but the relative heights of the barriers were appropriately scaled as follows. The difference between the heights of two alternative barriers for exiting a single source state was calculated as 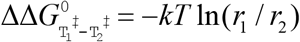, where *r*_1_ and *r*_2_ are the rates of exit along those two pathways. Similarly, the difference between the heights of a given barrier for two different constructs was calculated using the same equation, but with *r*_1_ and *r*_2_ reflecting the corresponding transition rates for constructs 1 and 2, respectively.

### In silico estimation of bond distances

*In silico* estimation of H-bond lengths was done by modeller (Webb and Sali, 2016) using the quasi-open CFTR structure (pdbid: 6msm) as template. For each mutant, 100 structures were simulated using standard parametrization. The generated models were assessed by the calculated DOPE (discrete optimized protein energy) score. In the best 50 models distances between the E1124 backbone carbonyl group and the two terminal nitrogens of the R117 side chain were calculated with Pymol. In the case of E1124Δ and E1126Δ the backbone carbonyl groups of both G1123 and G1125, or G1123 and E1124, respectively, were assessed, and the smaller distance was taken. Finally, for each mutant the mean and the standard error of the distance estimates obtained from the 50 models were calculated (Fig. 2 – Fig. Suppl. 2).

### Statistics

All values are given as mean±S.E.M., with the numbers of independent samples provided in each figure legend. Statistical significances of interaction free energies were calculated using Student’s t test, ΔΔ*G*_int_ is reported significantly different from zero for p<0.05.

## Acknowledgements

Supported by EU Horizon 2020 Research and Innovation Program grant 739593, MTA Lendület grant LP2017-14/2017, and Cystic Fibrosis Foundation Research Grant CSANAD21G0 to L.C. M.A.S. received support from the ÚNKP-20-3-I-SE-34 New National Excellence Program of the Ministry for Innovation and Technology from the source of the National Research, Development and Innovation Fund.

## Author contributions

L.C. designed the project, M.A.S. performed all experiments and analyzed the data, L.C. and M.A.S. interpreted the results and wrote the manuscript.

## Figure legends

**Figure 2 – Fig. Suppl. 1.**
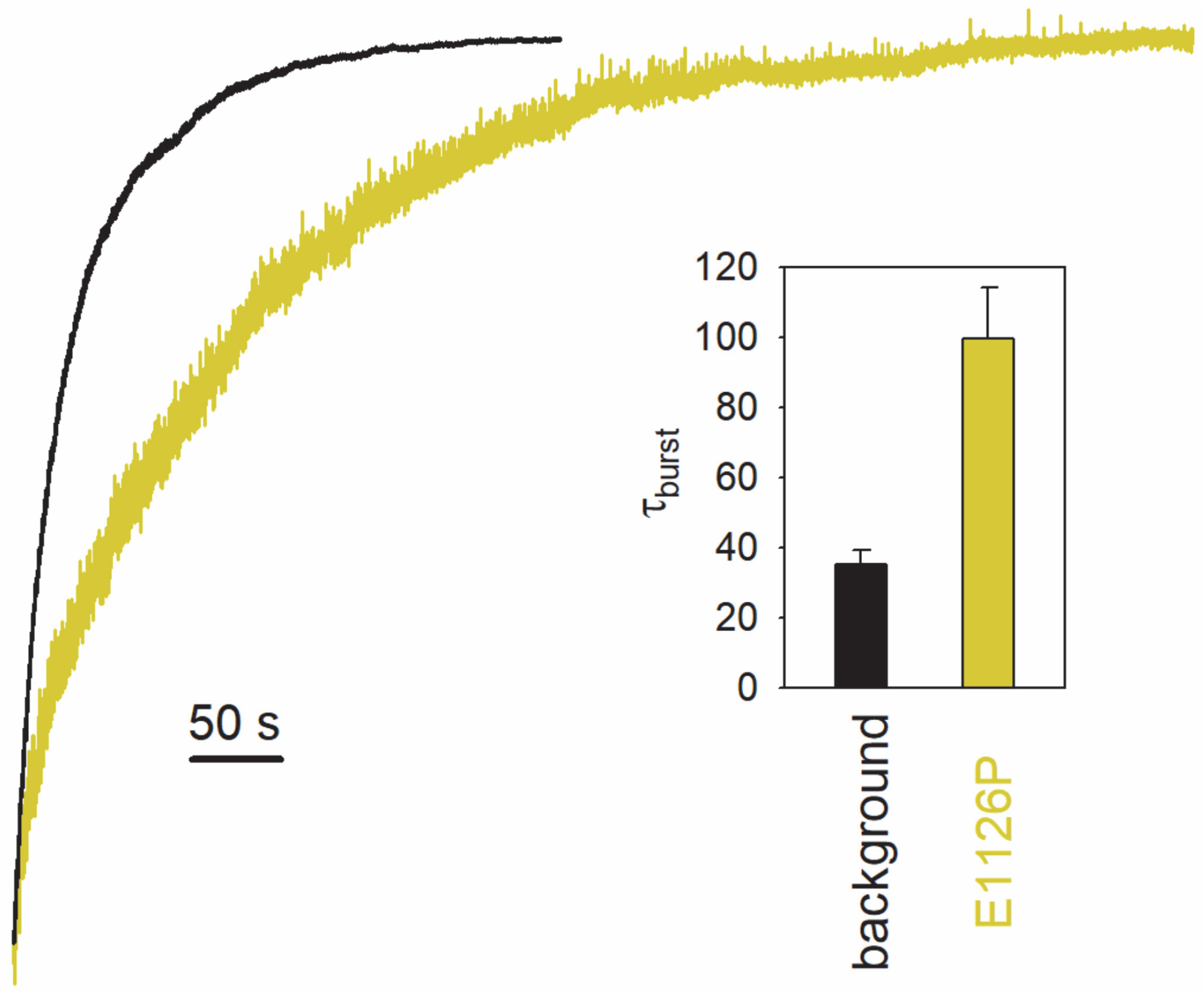
The E1126P mutation slows channel closure. Macroscopic current relaxations following ATP removal for E1371S (*black trace*) and E1371S-E1126P-CFTR (*yellow trace*). Inside-out patch currents were activated by exposure of pre-phosphorylated channels to 5 mM ATP. Current amplitudes are shown rescaled by their steady-state values. Membrane potential (V_m_) was -20 mV. (*Inset*) Relaxation time constants obtained by fits to single exponentials. Data are shown as mean±S.E.M. from 6 and 5 experiments, respectively.

**Figure 2 – Fig. Suppl. 2.**
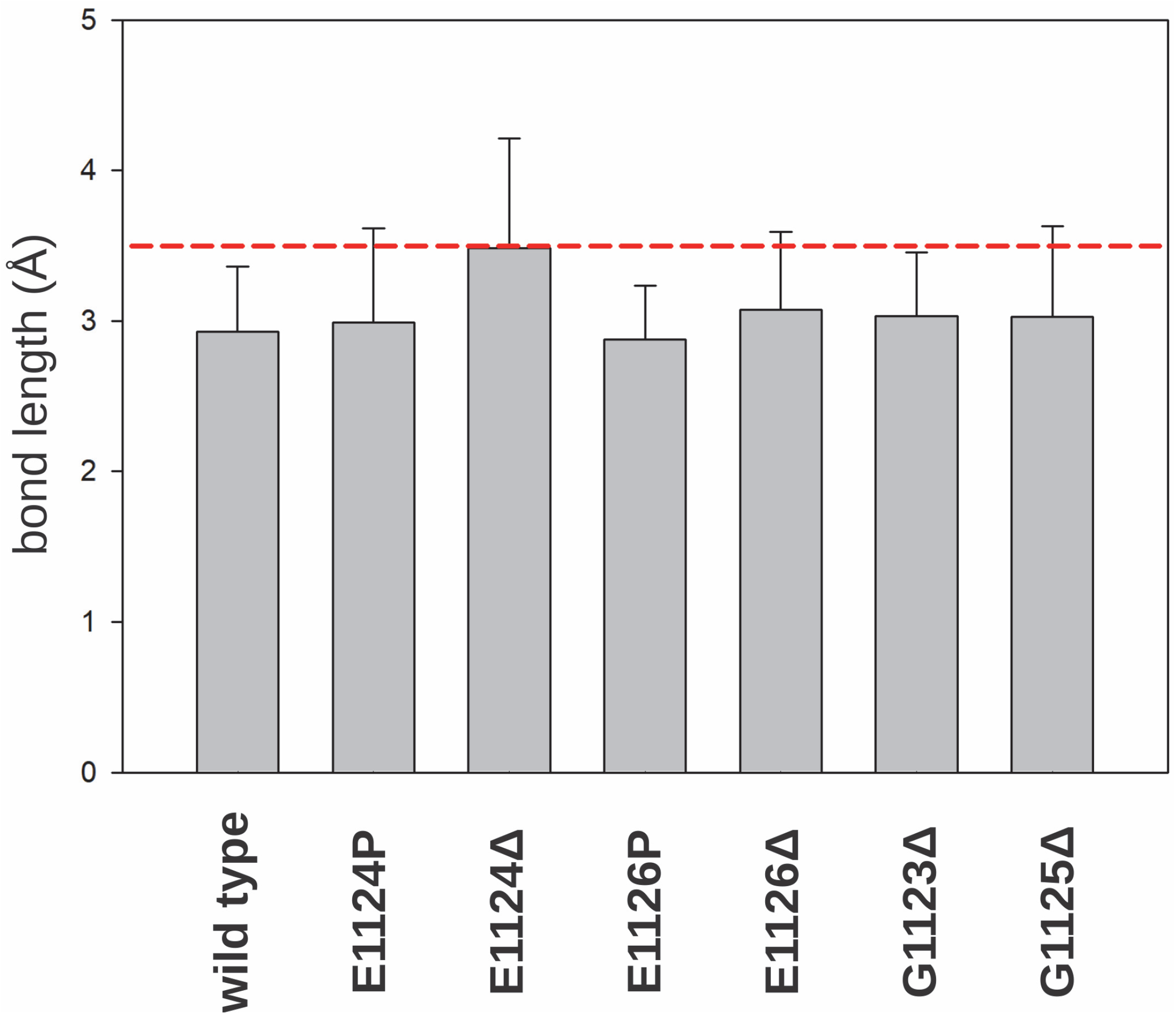
Estimated bond lengths of the target H-bond in various ECL6 mutants. *In silico* estimation of the smallest distance between the terminal nitrogen of the R117 side chain and the peptide carbonyl group of residue E1124 (or the nearest peptide carbonyl group) for various mutants with perturbations in ECL6, calculated based on the quasi-open human CFTR structure (6msm). See Materials and Methods for details. *Red line* denotes the cutoff distance for formation of an H-bond (Grabowski S.J., 2020).

**Figure 2 – Fig. Suppl. 3.**
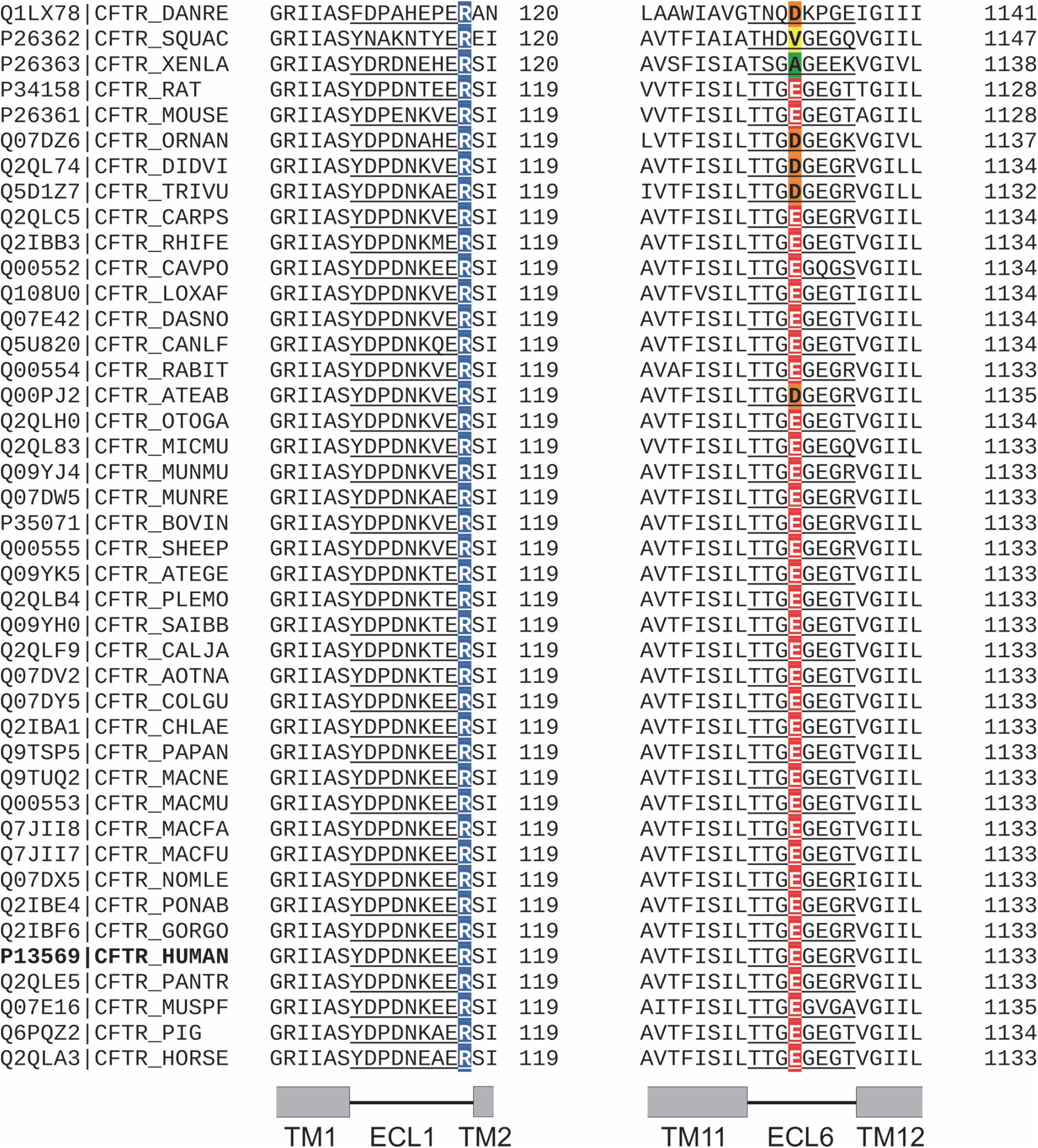
Sequence alignment of the ECL1 and ECL6 loops for CFTR orthologs. Segments of a multiple sequence alignment generated by Clustal Omega for CFTR sequences from >40 species. Positions corresponding to human CFTR positions 117 and 1124 are highlighted in *blue* and *red*, respectively. The former position is conserved, the latter is not. However, the lengths of both ECL1 and ECL6 are invariable (*underlined*).

**Figure 3 – Fig. Suppl. 1.**
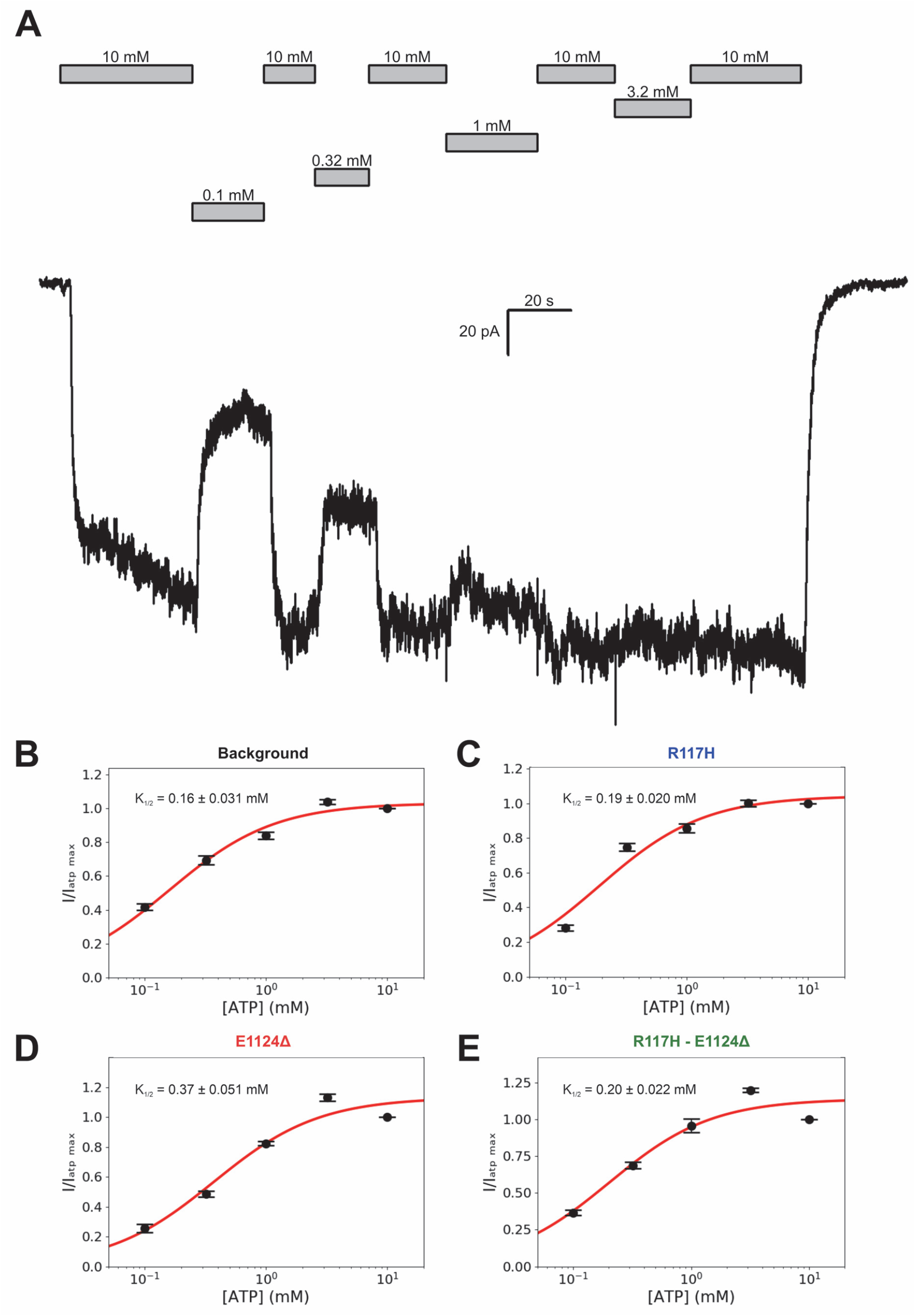
All tested constructs in the D1370N background are saturated by 5 mM ATP. *A*, Macroscopic current of pre-phosphorylated D1370N CFTR in response to exposure to various concentrations of ATP (*bars*). *B*-*E*, ATP dose-response curves of the four indicated constructs in the D1370N background. Relative currents were obtained by normalizing the steady-state currents in each test segment to the mean of the steady-state currents during bracketing applications of 10 mM ATP. Each symbols represents mean±S.E.M. from 5 experiments. Solid red curves are fits to the Michaelis-Menten equation, K_1/2_ values are plotted in the panels.

**Figure 3 – Fig. Suppl. 2.**
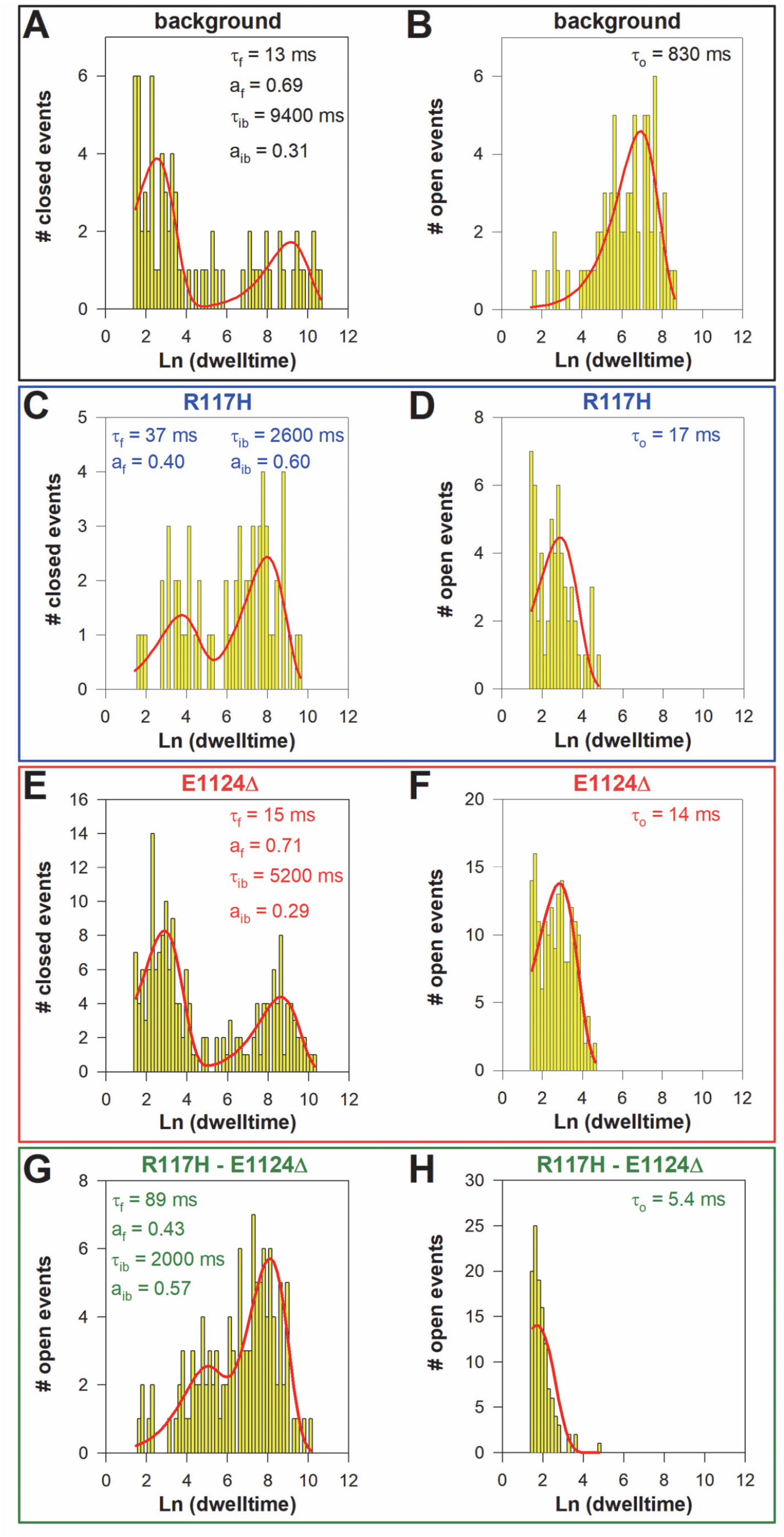
Steady-state dwell-time distributions of the target-site mutants in the D1370N background. *A*-*H*, Closed (*A, C, E, G*) and open (*B, D, F, H*) dwell-time histograms (Sigworth and Sine, 1987) constructed from the events list of single-channel records for the indicated CFTR mutants in the D1370N background; lower binning limit is 4 ms. *Solid red lines* are maximum likelihood fits of the dwell-times by the C_s_↔O↔C_f_ model (see Materials and Methods). Calculated time constants, and for the closed-time histograms the fractional amplitudes (a), are printed in the panels. The two time constants of the closed-time distributions correspond to τ_interburst_ and τ_flicker_, respectively, that of the open-time distribution to τ_open_.

**Figure 3 – Fig. Suppl. 3.**
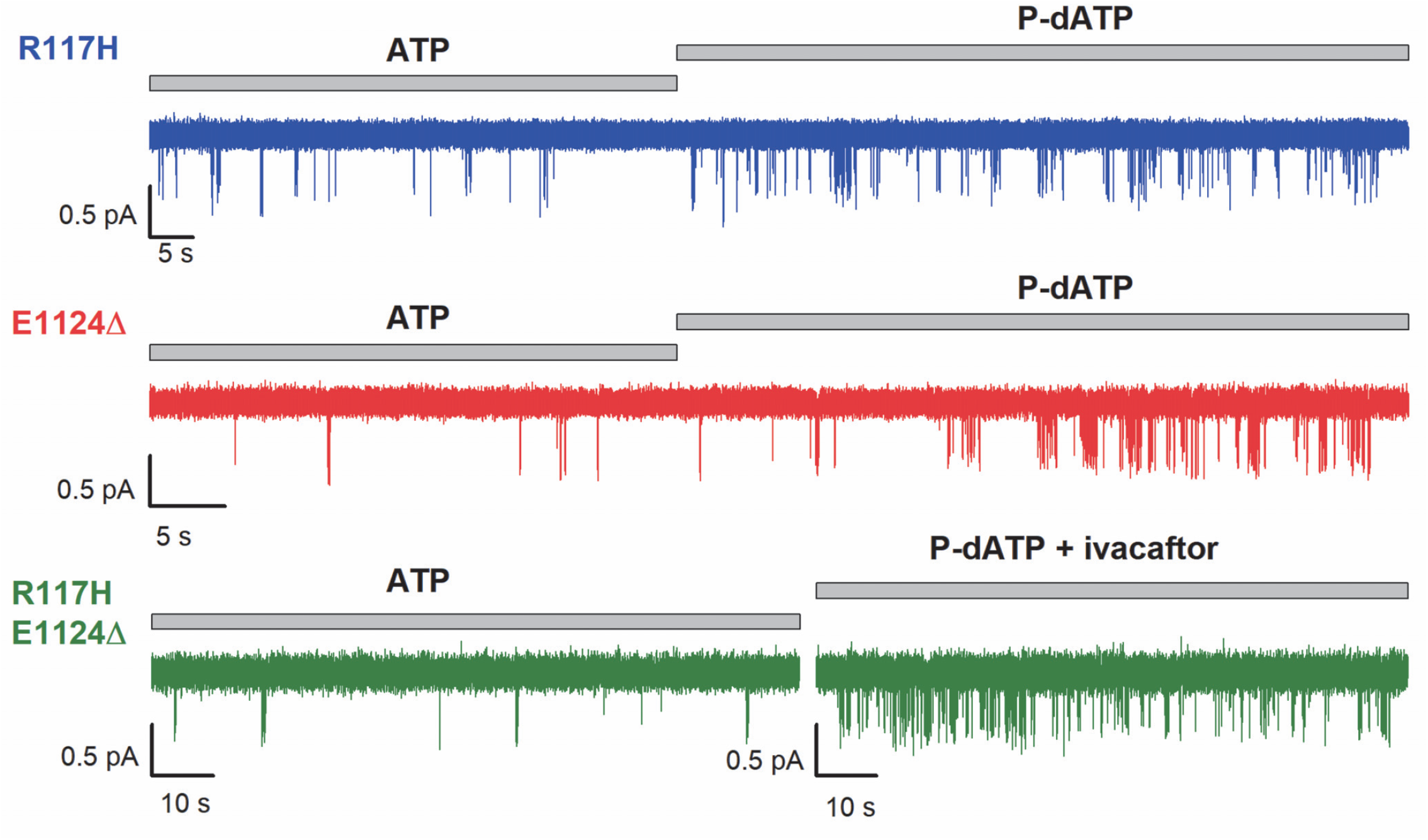
Stimulation by P-dATP or P-dATP+Vx-770 facilitates counting channels for low-P_o_ mutants. Single-channel recordings in 5 mM ATP of the three low-Po mutants in the D1370N background, and robust stimulation of P_o_ by 50 μM N^6^-(2-phenylethyl)-dATP (P-dATP), with or without 10 nM Vx-770 (ivacaftor). A statistical test based on comparison of the observed opening rate with the cumulative open time (Csanády et al., 2000) allowed high-confidence exclusion of the presence of a second active channel for each of the three patches shown (p<0.012, p<0.03, and p<0.003, respectively).

**Figure 5 – Fig. Suppl. 1.**
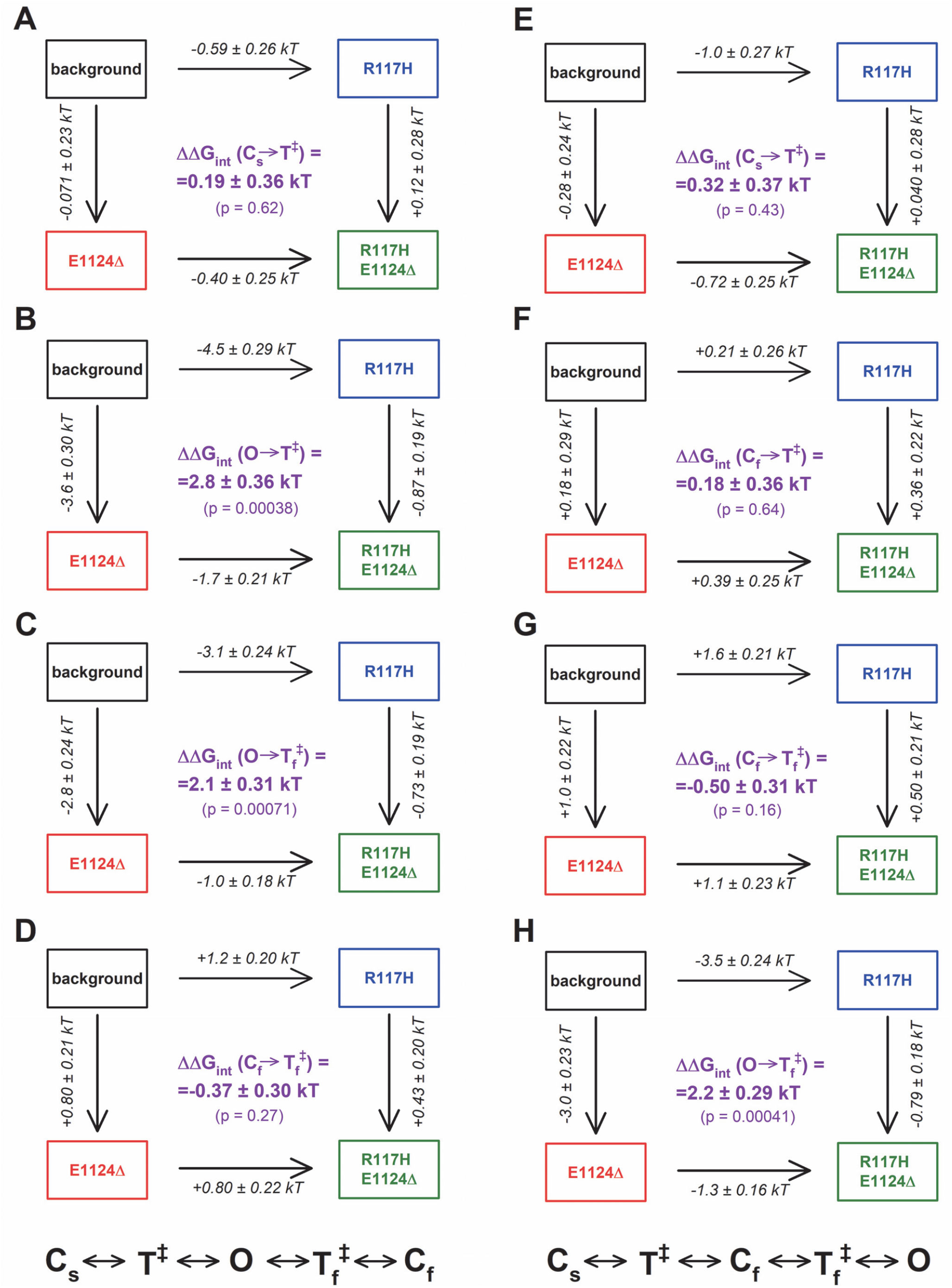
Mutant cycles built on the four microscopic transition rate constants for the two alternative three-state schemes. *A*-*H*, Thermodynamic mutant cycles showing mutation-induced changes in the heights of the free enthalpy barriers (ΔΔ*G*^0^) for the four microscopic transitions contained in the (*A*-*D*) C_s_↔O↔C_f_ or (*E*-*H*) C_s_↔C_f_↔O model, respectively (*numbers on arrows*; *k*, Boltzmann’s constant, *T*, absolute temperature). Each corner is represented by the mutations introduced into positions 117 and 1124 of D1370N-CFTR. ΔΔ*G*_int_ (*purple numbers*) are obtained as the difference between ΔΔ*G*^0^ values along two parallel sides of the cycle. Gating steps: *A*, C_s_→O; *B*, O→C_s_; *C*, O→C_f_; *D*, C_f_→O; *E*, C_s_→C_f_; *F*, C_f_→C_s_; *G*, C_f_→O; *H*, O→C_f_.

## References

Auerbach, A. 2007. How to turn the reaction coordinate into time. J Gen Physiol 130:543–546.

Bai, Y.H., M. Li, and T.C. Hwang. 2011. Structural basis for the channel function of a degraded ABC transporter, CFTR (ABCC7). J Gen Physiol 138:495–507.

Cai, Z., T.S. Scott-Ward, and D.N. Sheppard. 2003. Voltage-dependent gating of the cystic fibrosis transmembrane conductance regulator Cl-channel. J Gen Physiol 122:605–620.

Chen, J.H., W. Xu, and D.N. Sheppard. 2017. Altering intracellular pH reveals the kinetic basis of intraburst gating in the CFTR Cl-channel. J Physiol 595:1059–1076.

Colquhoun, D. and F.J. Sigworth. 1995. Fitting and statistical analysis of single-channel records. In Single channel recording. B.Sakmann and E.Neher, editors. Plenum Press, New York.

Csanády, L. 2000. Rapid kinetic analysis of multichannel records by a simultaneous fit to all dwell-time histograms. Biophys J 78:785–799.

Csanády, L. 2009. Application of rate-equilibrium free energy relationship analysis to nonequilibrium ion channel gating mechanisms. J Gen Physiol 134:129–136.

Csanády, L., K.W. Chan, D. Seto-Young, D.C. Kopsco, A.C. Nairn, and D.C. Gadsby. 2000. Severed channels probe regulation of gating of cystic fibrosis transmembrane conductance regulator by its cytoplasmic domains. J Gen Physiol 116:477–500.

Csanády, L., C. Mihályi, A. Szollosi, B. Torocsik, and P. Vergani. 2013. Conformational changes in the catalytically inactive nucleotide binding site of CFTR. J Gen Physiol 142:61–73.

Csanády, L., P. Vergani, and D.C. Gadsby. 2010. Strict coupling between CFTR’s catalytic cycle and gating of its Cl-ion pore revealed by distributions of open channel burst durations. Proc Natl Acad Sci U S A 107:1241–1246.

Csanády, L., P. Vergani, and D.C. Gadsby. 2019. Structure, gating, and regulation of the CFTR anion channel. Physiol Rev 99:707–738.

Cui, G.Y., K.S. Rahman, D.T. Infield, C. Kuang, C.Z. Prince, and N.A. McCarty. 2014. Three charged amino acids in extracellular loop 1 are involved in maintaining the outer pore architecture of CFTR. J Gen Physiol 144:159–179.

Davies, J.C., S.M. Moskowitz, C. Brown, A. Horsley, M.A. Mall, E.F. Mckone, B.J. Plant, D. Prais, B.W. Ramsey, J.L. Taylor-Cousar, E. Tullis, A. Uluer, C.M. McKee, S. Robertson, R.A. Shilling, C. Simard, F. Van Goor, D. Waltz, F. Xuan, T. Young, and S.M. Rowe. 2018. VX-659-Tezacaftor-Ivacaftor in Patients with Cystic Fibrosis and One or Two Phe508del Alleles. N Engl J Med 379:1599–1611.

De Boeck, K. and M.D. Amaral. 2016. Progress in therapies for cystic fibrosis. Lancet Respir Med 4:662–674.

Dean, M., M.B. White, J. Amos, B. Gerrard, C. Stewart, K.T. Khaw, and M. Leppert. 1990. Multiple mutations in highly conserved residues are found in mildly affected cystic fibrosis patients. Cell 61:863–870.

Grabowski S.J. 2020. Hydrogen Bond – Definitions, Criteria of Existence and Various Types. In Understanding Hydrogen Bonds: Theoretical and Experimental Views. Royal Society of Chemistry, 1–40.

Gunderson, K.L. and R.R. Kopito. 1995. Conformational states of CFTR associated with channel gating: the role ATP binding and hydrolysis. Cell 82:231–239.

Hammerle, M.M., A.A. Aleksandrov, and J.R. Riordan. 2001. Disease-associated mutations in the extracytoplasmic loops of cystic fibrosis transmembrane conductance regulator do not impede biosynthetic processing but impair chloride channel stability. J Biol Chem 276:14848–14854.

Hung, L.W., I.X. Wang, K. Nikaido, P.Q. Liu, G.F. Ames, and S.H. Kim. 1998. Crystal structure of the ATP-binding subunit of an ABC transporter. Nature 396:703–707.

Liu, F., Z. Zhang, L. Csanády, D.C. Gadsby, and J. Chen. 2017. Molecular Structure of the Human CFTR Ion Channel. Cell 169:85–95.

Mihályi, C., B. Torocsik, and L. Csanády. 2016. Obligate coupling of CFTR pore opening to tight nucleotide-binding domain dimerization. Elife 5. pii: e18164. doi: 10.7554/eLife.18164.:e18164.

O’Sullivan, B.P. and S.D. Freedman. 2009. Cystic fibrosis. Lancet 373:1891–1904.

Pless, S.A. and C.A. Ahern. 2013. Unnatural amino acids as probes of ligand-receptor interactions and their conformational consequences. Annu Rev Pharmacol Toxicol 53:211–229.

Rai, V., M. Gaur, S. Shukla, S. Shukla, S.V. Ambudkar, S.S. Komath, and R. Prasad. 2006. Conserved Asp327 of walker B motif in the N-terminal nucleotide binding domain (NBD-1) of Cdr1p of Candida albicans has acquired a new role in ATP hydrolysis. Biochemistry 45:14726–14739.

Ramsey, B.W., J. Davies, N.G. McElvaney, E. Tullis, S.C. Bell, P. Drevinek, M. Griese, E.F. Mckone, C.E. Wainwright, M.W. Konstan, R. Moss, F. Ratjen, I. Sermet-Gaudelus, S.M. Rowe, Q.M. Dong, S. Rodriguez, K. Yen, C. Ordonez, and J.S. Elborn. 2011. A CFTR Potentiator in Patients with Cystic Fibrosis and the G551D Mutation. N Engl J Med 365:1663–1672.

Riordan, J.R., J.M. Rommens, B. Kerem, N. Alon, R. Rozmahel, Z. Grzelczak, J. Zielenski, S. Lok, N. Plavsic, J.L. Chou, et al. 1989. Identification of the cystic fibrosis gene: cloning and characterization of complementary DNA. Science 245:1066–1073.

Sheppard, D.N., D.P. Rich, L.S. Ostedgaard, R.J. Gregory, A.E. Smith, and M.J. Welsh. 1993. Mutations in CFTR associated with mild-disease-form Cl-channels with altered pore properties. Nature 362:160–164.

Sigworth, F.J. and S.M. Sine. 1987. Data transformations for improved display and fitting of single-channel dwell time histograms. Biophys J 52:1047–1054.

Sorum, B., D. Czege, and L. Csanády. 2015. Timing of CFTR Pore Opening and Structure of Its Transition State. Cell 163:724–733.

Sorum, B., B. Torocsik, and L. Csanády. 2017. Asymmetry of movements in CFTR’s two ATP sites during pore opening serves their distinct functions. Elife 6. pii: e29013.

Van Goor, F., H. Yu, B. Burton, and B.J. Hoffman. 2014. Effect of ivacaftor on CFTR forms with missense mutations associated with defects in protein processing or function. J Cyst Fibros 13:29–36.

Vergani, P., S.W. Lockless, A.C. Nairn, and D.C. Gadsby. 2005. CFTR channel opening by ATP-driven tight dimerization of its nucleotide-binding domains. Nature 433:876–880.

Vergani, P., A.C. Nairn, and D.C. Gadsby. 2003. On the mechanism of MgATP-dependent gating of CFTR Cl-channels. J Gen Physiol 121:17–36.

Wang, W., B.C. Roessler, and K.L. Kirk. 2014. An Electrostatic Interaction at the Tetrahelix Bundle Promotes Phosphorylation-dependent Cystic Fibrosis Transmembrane Conductance Regulator (CFTR) Channel Opening. J Biol Chem 289:30364–30378.

Webb, B. and A. Sali. 2016. Comparative Protein Structure Modeling Using MODELLER. Curr Protoc Bioinformatics 54:5.

Wilschanski, M., J. Zielenski, D. Markiewicz, L.C. Tsui, M. Corey, H. Levison, and P.R. Durie. 1995. Correlation of sweat chloride concentration with classes of the cystic fibrosis transmembrane conductance regulator gene mutations. J Pediatr 127:705–710.

Winter, M.C., D.N. Sheppard, M.R. Carson, and M.J. Welsh. 1994. Effect of ATP concentration on CFTR Cl-channels: a kinetic analysis of channel regulation. Biophys J 66:1398–1403.

Yu, Y.C., Y. Sohma, and T.C. Hwang. 2016. On the mechanism of gating defects caused by the R117H mutation in cystic fibrosis transmembrane conductance regulator. J Physiol 594:3227–3244.

Zhang, J., Y.C. Yu, J.T. Yeh, and T.C. Hwang. 2018a. Functional characterization reveals that zebrafish CFTR prefers to occupy closed channel conformations. PLoS One 13:e0209862.

Zhang, Z., F. Liu, and J. Chen. 2017. Conformational Changes of CFTR upon Phosphorylation and ATP Binding. Cell 170:483–491.

Zhang, Z., F. Liu, and J. Chen. 2018b. Molecular structure of the ATP-bound, phosphorylated human CFTR. Proc Natl Acad Sci U S A.

Zhou, J.J., M. Fatehi, and P. Linsdell. 2008. Identification of positive charges situated at the outer mouth of the CFTR chloride channel pore. Pflugers Arch 457:351–360.

